# Action Potential Propagation and Synchronisation in Myelinated Axons

**DOI:** 10.1101/599746

**Authors:** Helmut Schmidt, Thomas R. Knösche

## Abstract

With the advent of advanced MRI techniques it has become possible to study axonal white matter non-invasively and in great detail. Measuring the various parameters of the long-range connections of the brain opens up the possibility to build and refine detailed models of large-scale neuronal activity. One particular challenge is to find a mathematical description of action potential propagation that is sufficiently simple, yet still biologically plausible to model signal transmission across entire axonal fibre bundles. We develop a mathematical framework in which we replace the Hodgkin-Huxley dynamics by a spike-diffuse-spike model with passive sub-threshold dynamics and explicit, threshold-activated ion channel currents. This allows us to study in detail the influence of the various model parameters on the action potential velocity and on the entrainment of action potentials between ephaptically coupled fibres without having to recur to numerical simulations. Specifically, we recover known results regarding the influence of axon diameter, node of Ranvier length and internode length on the velocity of action potentials. Additionally, we find that the velocity depends more strongly on the thickness of the myelin sheath than was suggested by previous theoretical studies. We further explain the slowing down and synchronisation of action potentials in ephaptically coupled fibres by their dynamic interaction. In summary, this study presents a solution to incorporate detailed axonal parameters into a whole-brain modelling framework.

**Author summary:** With more and more data becoming available on white-matter tracts, the need arises to develop modelling frameworks that incorporate these data at the whole-brain level. This requires the development of efficient mathematical schemes to study parameter dependencies that can then be matched with data, in particular the speed of action potentials that cause delays between brain regions. Here, we develop a method that describes the formation of action potentials by threshold activated currents, often referred to as spike-diffuse-spike modelling. A particular focus of our study is the dependence of the speed of action potentials on structural parameters. We find that the diameter of axons and the thickness of the myelin sheath have a strong influence on the speed, whereas the length of myelinated segments and node of Ranvier length have a lesser effect. In addition to examining single axons, we demonstrate that action potentials between nearby axons can synchronise and slow down their propagation speed.

## Introduction

Neurons communicate via chemical and electrical signals, and an integral part of this communication is the transmission of action potentials along their axons. The velocity of action potentials is crucial for the right timing in information processing and depends on the dynamics of ion channels studding the axon, but also on its geometrical properties. For instance, the velocity increases approximately linearly with the diameter of myelinated axons [1]. Myelin sheaths around axons are an evolutionary trait in most vertebrates and some invertebrates, which developed independently in several taxa [2]. The presence of a myelin sheath increases the velocity of action potentials by enabling saltatory conduction [3]. Long-term, activity-dependent changes in the myelination status of axons are related to learning [4]. The functional role of differentiated myelination is to regulate and synchronise signal transmission across different axonal fibres to enable cognitive function, sensory integration and motor skills [5]. White-matter architecture has also been found to affect the peak frequency of the alpha rhythm [6]. Axons and their supporting cells make up the white matter, which has, for a long time, only been accessible to histological studies [7, 8]. With the advent of advanced MRI techniques, some of the geometric parameters of axonal fibre bundles have become accessible to non-invasive methods. Techniques have been proposed to determine the orientation of fibre bundles in the white matter [9] as well as to estimate the distribution of axonal diameters [10], the packing density of axons in a fibre bundle [11, 12], and the ratio of the diameters of the axon and the myelin sheath (g-ratio) [13].

First quantitative studies were done by Hursh [14] who established the (approximately) linear relationship between action potential velocity and axonal radius in myelinated axons, and Tasaki [3] who first described saltatory conduction in myelinated axons. Seminal work on ion channel dynamics was later done by Hodgkin and Huxley, establishing the voltage-dependence of ion channel currents [15]. The general result of voltage-dependent gating has been confirmed in vertebrates [16], yet a recent result for mammals suggests that the gating dynamics of sodium channels is faster than described by the original Hodgkin-Huxley model, thereby enabling faster generation and transmission of action potentials [17]. In general, parameters determining channel dynamics differ widely across neuron types [18].

Seminal studies into signal propagation in myelinated axons using computational techniques were done by FitzHugh [19] and Goldman and Albus [20]. Goldman and Albus gave the first computational evidence for the linear increase of the conduction velocity with the radius of the axon, provided that the length of myelinated segments also increases linearly with the axonal radius. The linear relationship is supported by experimental evidence [21], although other studies suggest a slightly nonlinear relationship [22]. More recently, computational studies have investigated the role of the myelin sheath and the relationship between models of different complexity with experimental results [23]. One of the key findings here was that only a myelin sheath with finite capacitance and resistance reproduced experimental results for axonal conduction velocity. Other studies investigated the role of the width of the nodes of Ranvier on signal propagation [24, 25], or the effect of ephaptic coupling on signal propagation [26–33].

Most computational studies employ numerical schemes, i.e. they discretise the mathematical problem in space and time and use numerical integration methods to investigate the propagation of action potentials. One problem that arises here is that the spatial discretisation must be relatively coarse to ensure numerical stability, which can be remedied to some extent by advanced numerical methods and computational effort [34]. The other problem, however, cannot be remedied that easily: it is the lack of insight into how the model parameters influence the results, since there is a large number of parameters involved. A way to illustrate parameter dependencies in an efficient manner is to use analytical techniques all the while simplifying the model equations and extracting essential features. Studies that use analytical methods are few and far between [35–38]; yet it is also worth noting that from a mathematical perspective, myelinated axons are similar to spine-studded dendrites, in the sense that active units are coupled by passive leaky cables. An idea that we pick up from the latter is to simplify the ionic currents crossing the membrane [39, 40], there at dendritic spines mediated by neurotransmitters, here at nodes of Ranvier mediated by voltage-gated dynamics.

The goal of this article is to use analytical methods to study the influence of parameters controlling action potential generation, and geometric and electrophysiological parameters of the myelinated axon, on the speed of action potentials. The main focus here is on parameters determining the axonal structure. This will be achieved by replacing the Hodgkin-Huxley dynamics with a spike-diffuse-spike model for action potential generation, i.e. ion currents are released at nodes of Ranvier when the membrane potential reaches a certain threshold. These ion channel currents are considered voltage-independent, but we investigate different forms of currents, ranging from instantaneous currents to currents that incorporate time delays. We also investigate ion currents that closely resemble sodium currents measured experimentally. Our aim is to derive closed-form solutions for the membrane potential along an axon, which yields the relationship of action potential velocity with model parameters.

The specific questions we seek to answer here are the following. First, we query how physiological parameters can be incorporated into our mathematical framework, especially parameters that control the dynamics of the ionic currents. We test if parameters from the literature yield physiologically plausible results for the shape and amplitude of action potentials, and test how the ionic currents from multiple nearby nodes of Ranvier contribute to the formation of action potentials. Secondly, we ask how geometric parameters of an axon affect the transmission speed in a single axon, and how sensitive the transmission speed is to changes in these parameters. We seek to reproduce known results from the literature, such as the dependence of the velocity on axon diameter. We also explore other dependencies, such as on the g-ratio, and other microscopic structural parameters resulting from myelination. We compare the results of our spike-diffuse-spike model with the results from a detailed biophysical model recently used to study the effect of node and internode length on action potential velocity [24]. Thirdly, we investigate how ephaptic coupling affects the transmission speed of action potentials, and what the conditions are for action potentials to synchronise. In particular, we examine how restricted extra-axonal space leads to coupling between two identical axons, and how action potentials travelling through the coupled axons interact.

## Results

For the mathematical treatment of action potential propagation along myelinated axons, we consider active elements periodically placed on an infinitely long cable. The latter represents the myelinated axon and is appropriately described as leaky cable, whereas the active elements represent the nodes of Ranvier. In mathematical terms, the governing equation is an inhomogeneous cable equation, which describes the membrane potential *V* (*x, t*) of a leaky cable in space *x* (scalar, longitudinal to the cable) and time *t* in response to input currents:

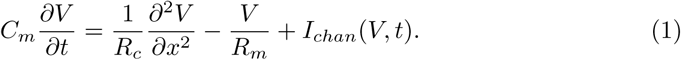

Here, *C*_*m*_ and *R*_*m*_ are the (radial) capacitance and resistance of a myelinated fibre, and *R*_*c*_ is its axial resistance. The term *I*_*chan*_ represents the ion channel currents triggered at nodes of Ranvier. The cable equation (1) can be reformulated into

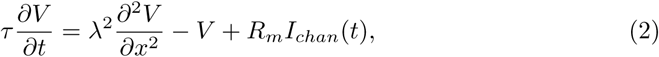

by multiplying both sides of (1) with *R*_*m*_. The time constant *τ* and the cable constant *λ* are parameters determined by the electrophysiological properties of myelin. We choose these parameters in accordance with experimental results and keep them fixed throughout our analysis, see the Methods section for details.

The input currents generated by the ion channel dynamics at the nodes of Ranvier is commonly described by a Hodgkin-Huxley framework. However, the Hodgkin-Huxley equations are a challenge to solve analytically, and in order to proceed with our mathematical treatment we opt for a simplified description using threshold-activated currents with standardised current profiles. We analyse different current profiles, ranging from delta-spikes to combinations of exponentials which give a good approximation of the ion currents observed experimentally. We solve the cable equation for these currents analytically which yields the dynamics of the membrane potential describing the resulting depolarisation / hyperpolarisation along the axon. The linearity of the cable equation in *V* allows us to describe the response to multiple input currents by the superposition of solutions for single currents. A sketch of the framework is shown in Fig. 1.

**Fig 1.**
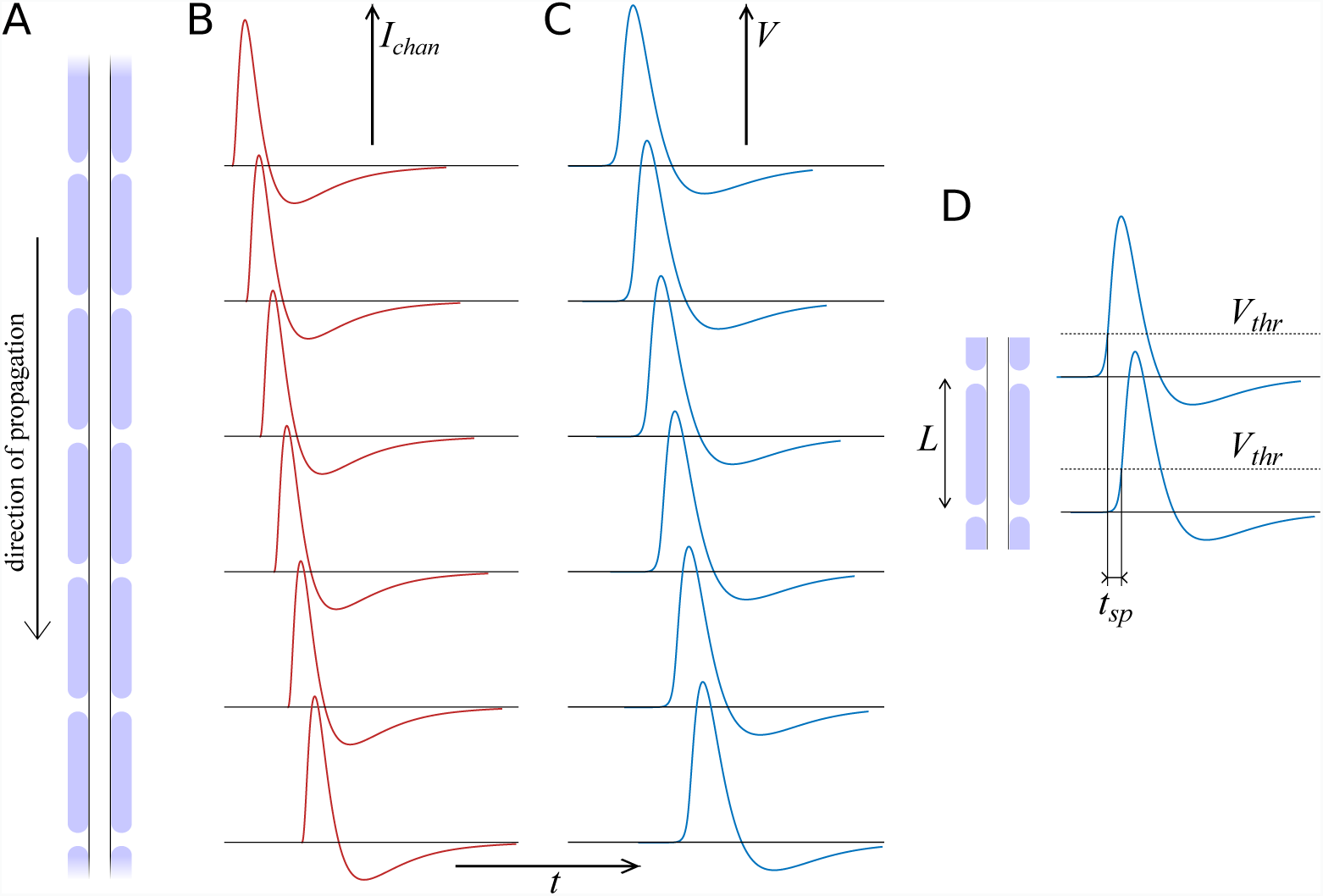
Action potential propagation in a myelinated axon. **A**: The axon is made of myelinated segments (internodes), with the nodes of Ranvier forming periodic gaps in the myelin sheath. **B**: The nodes of Ranvier constitute active sites at which threshold-triggered ion channel currents are released. **C**: The currents entering nearby nodes of Ranvier determine the membrane potential at each node, thus forming an action potential. **D** The velocity of an action potential is determined by the distance *L* between two consecutive nodes, and the time difference *t*_*sp*_ it takes to reach a given threshold value.

### Ion channel dynamics

The classical Hodgkin-Huxley model is described by a set of nonlinear equations which need to be solved numerically. Over the years, it has seen several modifications and improvements such as the one by Frankenhaeuser and Huxley [16], or the incorporation of additional ion currents [41] given the multitude of ion channel types [42, 43]. Also, attempts were made to provide better fits by modifying the exponents of the gating variables [44]. In essence, it is difficult to determine what is the ‘right’ Hodgkin-Huxley model for specific neuron types. For this reason, it seems prudent to go into the opposite direction and to try to simplify the description of the ion channel dynamics.

Two important contributions into this direction are the one by Fitzhugh [45, 46] and Nagumo [47], and the one by Morris and Lecar [48]. They provide a framework in which the slow and the fast variables are lumped and thus yield a two-dimensional reduction of the Hodgkin-Huxley model. The ion currents here are still voltage-dependent.

A crucial simplification towards analytically treatable models is the separation of sub-threshold dynamics and spike generation in integrate-and-fire models [49, 50]. For instance, in the leaky integrate-and-fire model and the quadratic integrate-and-fire model, the time-to-spike can be computed analytically, given initial conditions and a threshold value for the membrane potential. The ion currents are then often modelled as delta-spikes since the ion dynamics is fast in comparison to the (dendritic and somatic) membrane dynamics. The spatial extension of the leaky integrate-and-fire model is the spike-diffuse-spike model, in which activity spreads via passive cables.

Here, we consider four forms of channel current models. All of these have in common that the ion current is initiated after the membrane potential has crossed a threshold *V*_*thr*_, and has a predetermined profile. We denote the four scenarios by the letters A, B, C, and D. In scenario A, the ion channel current is released immediately and instantaneously, i.e.

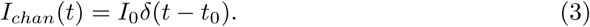

Here, *I*_0_ denotes the overall ion current, *t*_0_ denotes the time when the membrane potential crosses the threshold, and *δ*(⋅) is the delta-distribution, or Dirac’s delta. In scenario B, the ion current is also released instantaneously, but with a delay ∆:

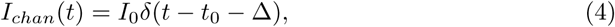

In scenario C, the ionic current is exponential:

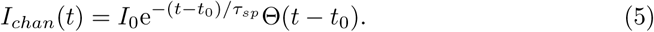

Here, *τ*_*sp*_ is the decay time, and Θ(⋅) is the Heaviside step function. With scenario D we aim to approximate the ion currents as measured in mammals such as the rabbit [51] and in the rat [52], which can be described by a superposition of exponential currents:

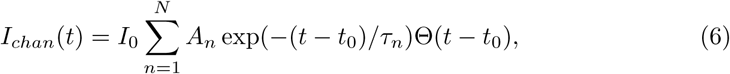

A sketch of all these scenarios is shown in Figure 2, alongside typical depolarisation curves of the membrane potential.

**Fig 2.**
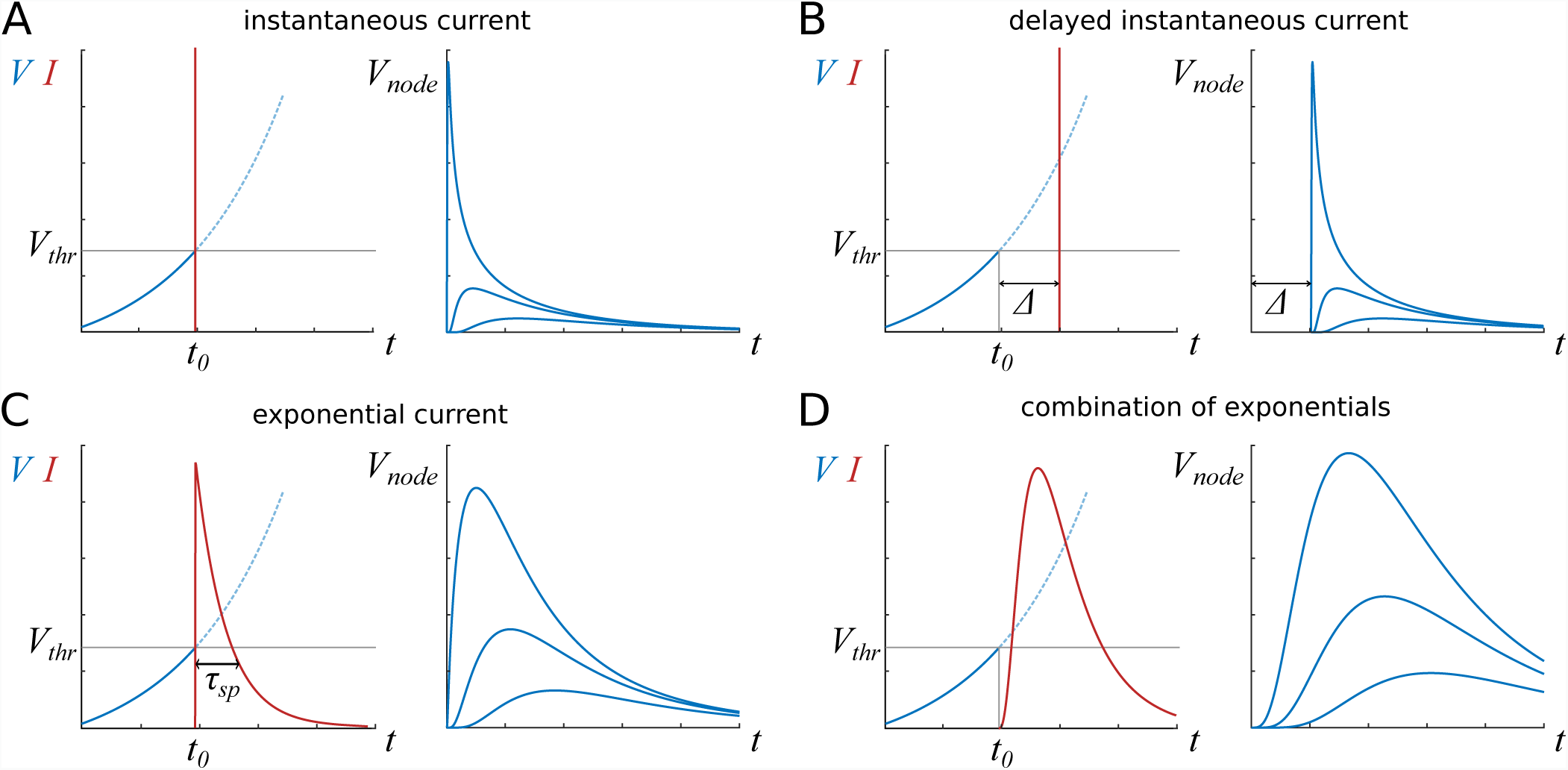
Sketch of ion channel currents considered here, with representative profiles of membrane potential in nearby nodes. After the membrane potential *V* reaches the threshold value *V*_*thr*_, the current *I* is released. **A**: The instantaneous current is described by a delta-peak at *t*_0_, when the threshold value is reached. **B**: The simplest way to accommodate delays or refractoriness is to introduce a refractory period ∆, after which the instantaneous current is released. **C**: Exponential current with characteristic time scale *τ*_*sp*_. **D:** A combination of exponential currents describes a realistic current profile.

### Current influx and separation

According to Kirchhoff’s first law, the channel current that flows into the axon, *I*_*chan*_(*t*) is counter-balanced by currents flowing axially both ways along the axon, *I*_*cable*_(*t*), and a radial current that flows back out across the membrane of the node, *I*_*node*_, see Fig. 3A for a graphical representation. The ratio of currents that pass along the cable and back across the nodal membrane is determined by the respective resistances:

**Fig 3.**
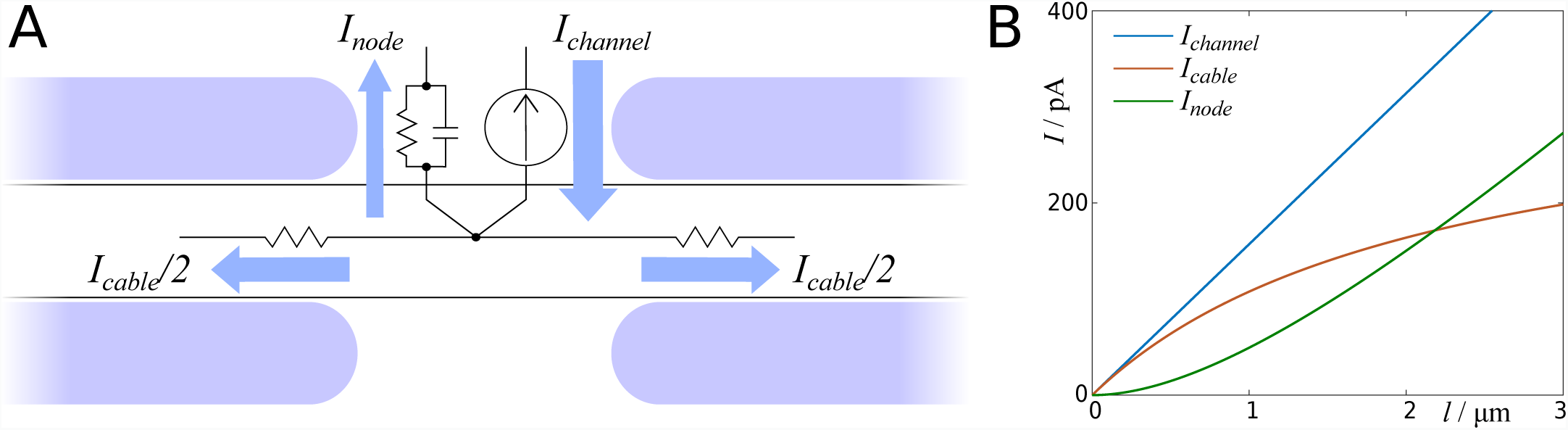
Channel currents divide into a current entering the axon and a current flowing back across the node of Ranvier. **A**: Sketch of currents entering and leaving a node of Ranvier. **B**: Plot of currents as function of node length. Since we assume constant channel density, the channel current increases linearly with the node length.

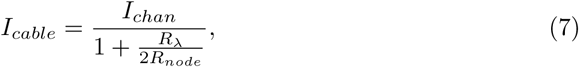

with *R*_*λ*_ = *R*_*m*_/*λ*. Throughout the manuscript, the ratio between *I*_*cable*_ and *I*_*chan*_ is expressed by *β*:

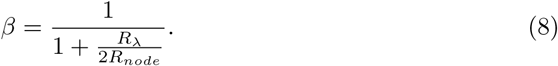

Based on experimental findings, we assume that the channel density is constant [52], which implies that the total channel current increases linearly with the node length. This is counterbalanced by the fact that the inverse of the resistance of a node, 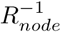, also increases linearly with its length. At large node lengths, the current that enters the axon saturates, see Fig. 3B. We will examine further below how the node length influences the propagation speed.

### Influence of nearby nodes

During the propagation of an action potential, ion channel currents are released at multiple nearby nodes that affect the shape and amplitude of the action potential. Because of the linear nature of the cable equation, the effect of multiple input currents can be described by linear superposition:

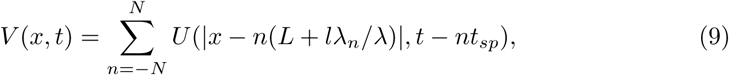

where *U* describes the depolarisation due to the current at a single node with index *n*. The internode length *L*, node length *l*, cable constant *λ* and cable constant at a node *λ*_*n*_ determine the electrotonic distance between nodes. Node indices *n* are chosen such that the node with *n* = 0 is centred at *x* = 0. Nodes with negative *n* are the ones the action potential has travelled past, and nodes with positive *n* are the ones the action potential will travel into. Although we consider infinitely long axons, we cut off the sum at *n* = *−N* and *n* = *N* for computational feasibility, with *N* = 10^3^. The action potential is not only shaped by the currents from preceding nodes, but also by currents from subsequent nodes that travel back along the axon. Due to the periodic nature of saltatory conduction, the time difference between any two consecutive nodes is assumed to be the same unknown parameter *t*_*sp*_.

The effect of distant nodes is dampened by the fact that in addition to passing along myelinated segments, currents from distant sources also pass by unmyelinated nodes, and therefore further lose amplitude. If nodes are relatively short, the current outflux can be regarded as instantaneous across the node as compared to changes in the current, and the total electrotonic distance between two consecutive nodes (measured in units of *λ*) is then given by *L* + *lλ/λ*_*n*_, which is already included in Eq. (9). Here, *λ*_*n*_ denotes the cable constant at a node. Eq. (9) describes the temporal evolution of an action potential in a specific location *x*. In Fig. 4 we dissect an action potential using scenario D for the ion channel model, by colour-coding the depolarisation due to individual nodes. It is apparent that the action potential propagation is a collective process with each node regenerating the action potential by a small fraction.

**Fig 4.**
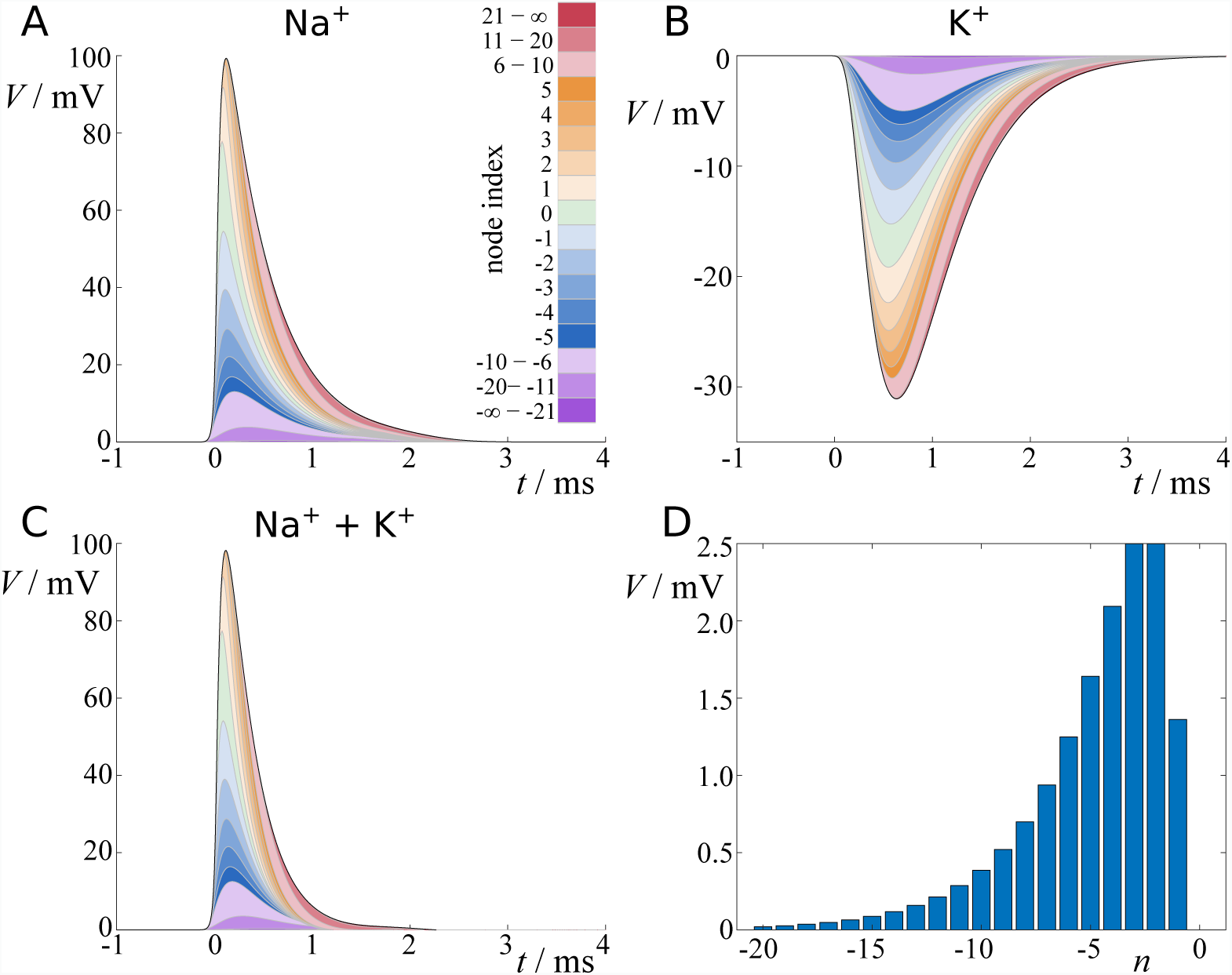
Contribution of ion currents from nearby nodes to action potential profile. **A**: Sodium currents contributing to action potential, and **B**: same for potassium. Depolarising effect is color-coded by node index, larger indices are lumped. Total effect is indicated by black line. **C**: Action potential composed of both currents. **D**: Contribution of sodium currents to reaching threshold value. Standard parameters are used here (Table 1 in Methods).

### Velocity of action potentials

We now consider the node at *x* = 0 (*n* = 0) to reach the firing threshold *V*_*thr*_ at *t* = 0. The relationship between the firing threshold *V*_*thr*_ and the time-to-spike *t*_*sp*_ is then given by

**Table 1.** List of model parameters used in this manuscript.

| Parameter | standard values | fitted values (AC model) |
| --- | --- | --- |
| $C_m$ | $3.6/\ln(1/g)pFcm^{-1}$ | N.A. |
| $R_m$ | $130M\Omega cm \ln(1/g)$ | $130M\Omega cm \ln(1/g)$ |
| $R_c$ | $140\Omega cm/d^2$ | N.A. |
| $\tau$ | $0.47ms$ | $1.45ms$ |
| $\tau_n$ | $33\mu s$ | $20\mu s$ |
| $\lambda$ | $9.65 \times 10^2 d \sqrt{\ln g^{-1}}$ | $12 \times 10^2 d \sqrt{\ln g^{-1}}$ |
| $\lambda_n$ | $38.9 \sqrt{d/\mu m} \mu m$ | $48.1 \sqrt{d/\mu m} \mu m$ |
| $d$ | $1\mu m$ | $0.73\mu m$ |
| $g$ | $0.6$ | $0.81$ |
| $l$ | $1\mu m$ | $1\mu m$ |
| $L$ | $100d$ | $100d$ |
| $\tau_m$ | $20\mu s$ | $70\mu s$ |
| $\tau_h$ | $40\mu s$ | $160\mu s$ |
| $\tau_n$ | $150\mu s$ | $150\mu s$ |
| $\tau_k$ | $300\mu s$ | $300\mu s$ |
| $I_0$ | $50pA/\mu m^2$ | $200pA/\mu m^2$ |
| $V_{thr}$ | $15mV$ | $4mV$ |

Unless explicitly stated, we use the parameters presented in this list. For most figures we use the standard parameters, and where stated we use parameters fitted to the Arancibia-Carcamo cortex model. The fitting procedure is described in the subsection ‘Fitting parameters to biophysical model’.

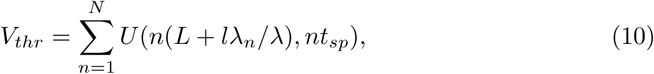

where we have changed the sign of the summation index, i.e. *−n → n*. The choice of *x* = 0 and *t* = 0 is without loss of generality. Eq. (10) is an implicit equation for *t*_*sp*_, which we solve here numerically using Newton’s method. The velocity of an action potential is then given by the physical distance between two consecutive nodes, *L* + *l*, and *t*_*sp*_:

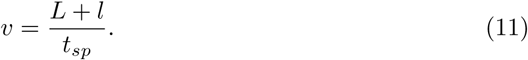

Here we still assume that the activation process at a node is uniform across its entire length. Since a node represents a short section of unmyelinated axon, we estimate the action potential velocity within a node by the action potential velocity in an unmyelinated axon, *v*_*n*_ (see Methods section). The resulting velocity then reads

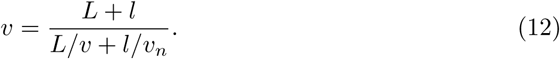

We use Eq. (12) throughout the manuscript.

### Analytical solutions

In mathematical terms, the depolarisation *U* resulting from the ion channel current at a single site, is a convolution of the current entering the cable with the Green’s function of the homogeneous cable equation *G*(*t*), which describes the propagation of depolarisation along the myelinated segment:

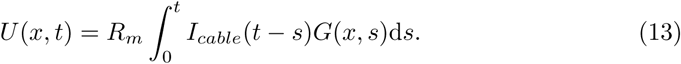

Here, *x* denotes the distance between the site where the current is injected and the site where the membrane potential is recorded. In the following we present the analytical solutions for all the current types.

#### Scenario A - fast current

Since the fast current is described by a delta function, the convolution integral turns into the Green’s function up to a prefactor:

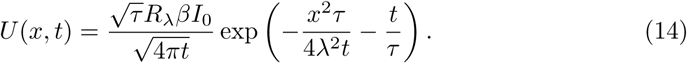

Here, *I*_0_ = 6.6*pA/µm*^2^ is the amplitude of the input current, *R*_*λ*_ = *R*_*m*_/*λ*, and *β* is the ratio between the current entering the cable and the channel current, as given by Eq (8). *I*_0_ is chosen such that the amplitude of an action potential is approximately 100*mV*, with all the other parameters chosen as for scenario D with standard parameters, see Methods section.

Inserting Eq. (14) into Eq. (9), we obtain the spatio-temporal evolution of an action potential for this scenario:

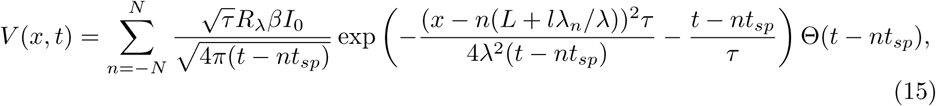

with Θ being the Heaviside step function to ensure causality. The threshold condition then reads

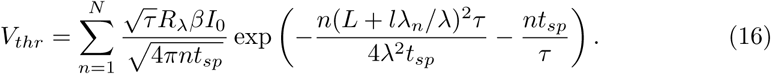

Although this is the simplest scenario, it is not obvious how to invert the r.h.s. of Eq. (16) to obtain an explicit expression for *t*_*sp*_. In the Methods section we present a linearisation approach, but it is convenient to solve Eq. (16) numerically using Newton’s method.

#### Scenario B - delayed fast current

The membrane dynamics in scenario B is exactly the same as in scenario A, except for an additional offset ∆:

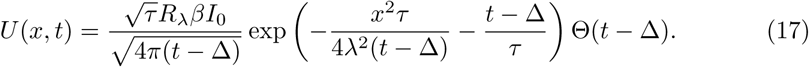

The spatio-temporal evolution of an action potential is now given by

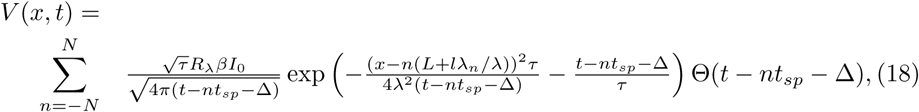

and the threshold condition reads

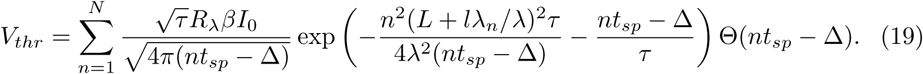

Because multiple nodes contribute to the depolarisation, it is possible to find *t*_*sp*_ *<* ∆.

#### Scenario C - exponential current

Here we have to solve the convolution integral of the cable equation with an exponential function, which yields

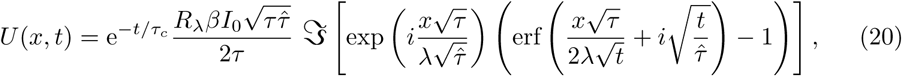

with 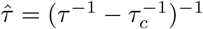, ℑ representing the imaginary part of the argument, and erf being the error function. In the Methods section we show how to obtain this solution. Eq. (20) thus represents solutions for ion currents with instantaneous onset and exponential decay. Hence, the spatio-temporal evolution of an action potential is expressed by

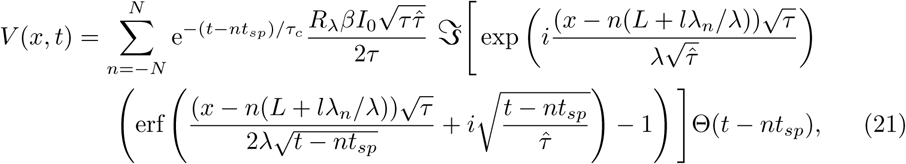

and the threshold condition to determine *t*_*sp*_ is

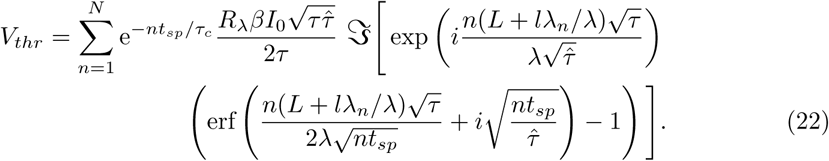

#### Scenario D - combination of exponentials

The linearity of the cable equation allows us to recur to the solution for scenario C to describe the response to currents described by multiple exponentials. Denoting the solution for one exponential input current with time constant *τ*_*s*_ by

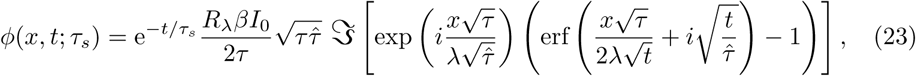

we express the solution to *M* superimposed exponential currents by

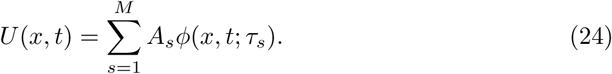

We use this formulation to describe both sodium currents and potassium currents with rising and falling phase. The sodium current is expressed as follows:

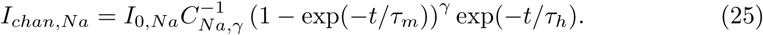

For simplicity, we focus on the case *γ* = 1, i.e. the biexponential case. Increasing *γ* would result in increased initial delays, and therefore lower propagation velocities. The parameter *γ* also affects the normalisation constant *C*_*Na,γ*_, which ensures that the maximum of *I*_*chan,Na*_ is *I*_0,*Na*_. The potassium current is modeled as

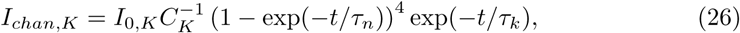

throughout the manuscript. In the Methods section we describe how to compute the normalisation constants *C*_*Na,γ*_ and *C*_*K*_, and how to convert Eq (25) and Eq (26) into a sum of exponentials. Hence, the spatio-temporal evolution of an action potential is expressed by

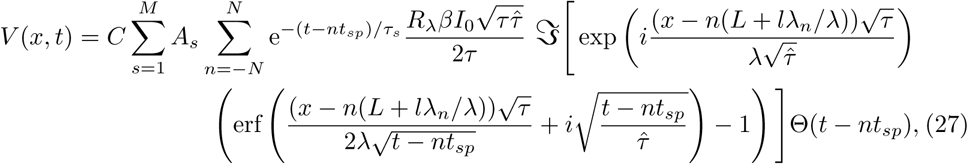

with 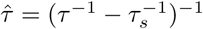, and *C* is the problem-specific normalisation constant. The threshold condition to determine *t*_*sp*_ is

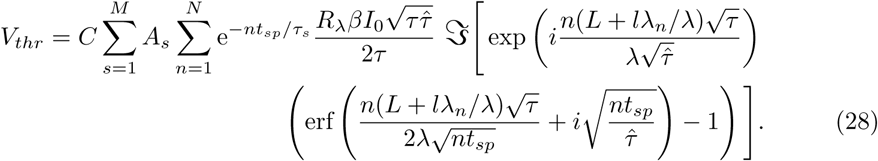

Anticipating results from the next subsection, we found that scenarios A and C yield velocities that are too fast compared with experimental results. Scenario B allows to adjust the propagation speed by tuning the parameter ∆, yet the shape of the action potential is only determined by the parameters from the cable equation, and thus cannot be adjusted to match experimental results. As it is the most realistic and most flexible model for ion channel currents, we decided to select scenario D to study the sensitivity of the propagation speed to structural parameters.

### Sensitivity to parameters

#### Axon diameter

There is a wide consensus that the propagation velocity in myelinated axons is proportional to the axon diameter. This is mostly due to the fact that both the internode length as well as the electrotonic length constant increase with the diameter. One quantity that does not scale linearly with the axonal diameter is the node length, which determines the amount of current that flows into the axon, as well as setting a correction term for the physical and electrotonic distance between two nodes. We find that the latter introduces a slight nonlinearity at small diameters, although at larger diameters the linear relationship is well preserved, see Fig. 5A.

**Fig 5.**
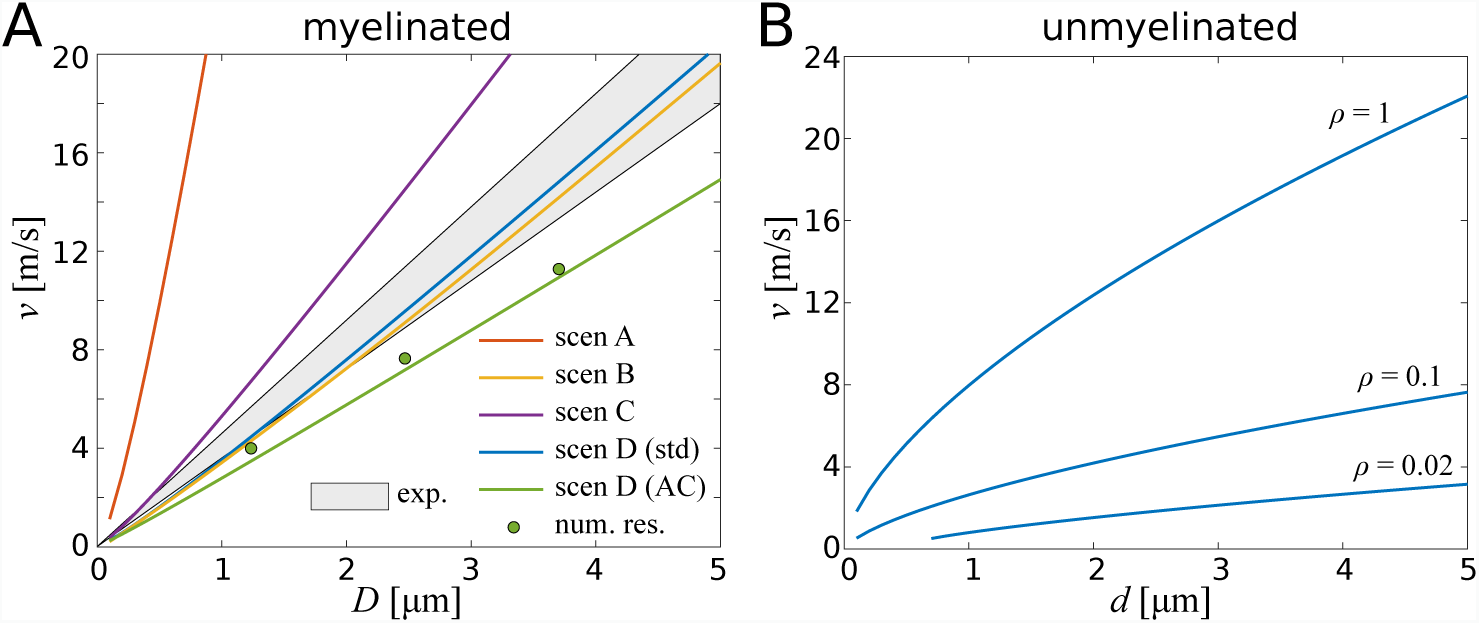
Propagation velocity as function of fibre diameter and axon diameter. **A**: In myelinated axons, the relationship between velocity and fibre diameter is nearly linear, with a slightly supralinear relationship at small diameters. Here we compare the different scenarios with experimental results (grey-shaded area). **B**: In unmyelinated axons, the propagation speed increases approximately with the square root of the axon diameter. Here, *ρ* indicates the relative ion channel density compared with a node of Ranvier. Decreasing the ion channel density results in slower action potential propagation.

In Fig. 5A we compare the four ion channel scenarios with experimental results obtained by Boyd and Kalu [53]. Scenario A (instantaneous ion channel current) yields velocities that are about one order of magnitude larger than the experimental results. This suggests that the main bottleneck for faster action potential propagation is indeed ion channel dynamics and their associated delays. Introducing a hard delay with scenario B, we find that we can reproduce the experimentally observed range of velocities. With scenarios C and D we introduce temporally distributed ion channel dynamics. The instantaneous onset and exponential decay of scenario C yields velocities that are slightly faster than experimental results.

In scenario D we explore two sets of parameters. The first set of parameters is obtained by using electrophysiological parameters found in the literature. As it is not obvious how to choose the time constants governing the temporal profile of the ion channel currents, we decided to choose them such that the shape of action potentials of our spike-diffuse-spike model match the shape of action potentials of the biophysical model used by Arancibia-Carcamo *et al.* [24]. The velocities obtained with this set of parameters fall within the range of experimental results. The second set of parameters is obtained by fitting the model parameters to data generated by the same biophysical model (see Methods). The latter yields velocities slightly below the experimental range, but it matches well the results from the biophysical model.

The present framework also enables us to study unmyelinated axons, in which case the current influx must be adapted, in addition to the physical and electrotonic distance between two neighbouring nodes, which is *l* and *l/λ*_*n*_, respectively. Since *λ*_*n*_ is proportional to 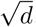, the resulting velocity is also to be expected to scale with 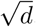, see Fig. 5B. Making the assumption that the membrane conductivity scales linearly with the ion channel density *ρ* (*ρ* is measured relative to the ion channel density of a node), the time constant of the unmyelinated axon scales with *τ* = *τ*_*n*_/*ρ*, and the cable constant scales with 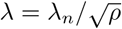. We study different ion channel densities, beginning with the same density as in nodes in the myelinated axon, and then reducing the density to 10% and 2% of the original density. We find that reducing the ion channel density also decreases the propagation velocity. For *ρ* = 1 we find that the propagation velocity is considerably faster than in myelinated axons at small diameters.

#### Node and internode length

Two geometric parameters that are not readily accessible to non-invasive MR-techniques are the length of the nodes of Ranvier, and the length of internodes. Here we examine the effect of the node and internode length on the speed of action potentials. We assume that the channel density in a node is constant, which is in agreement with experimental results [52]. The channel current that enters the node is proportional to its length, yet the increase of the node length also means that more of this current flows back across the node rather than entering the internodes. Another effect of the node length is the additional drop-off of the amplitude of axonal currents. Node lengths are known to vary between 1*µ*m and 3*µ*m [24].

The length of internodes is known to increase with the fibre diameter [21, 22]. This increase can be understood in light of the fact that the cable constant *λ* is proportional to the fibre diameter, and therefore increasing the internode length ensures that the ratio *L/λ* remains at a suitable point for signal transmission.

We restrict the analysis to the activation by sodium currents, since potassium currents are slow and only play a minor role in the initial depolarisation to threshold value. The results are shown graphically for scenario D with standard parameters in Fig. 6A, and for parameters fitted to the biophysical model by Arancibia-Carcamo *et al.* [24] in Fig. 6B. Changing the threshold value did have a small effect on the maximum velocity, but did not change the relative dependence on the other parameters.

**Fig 6.**
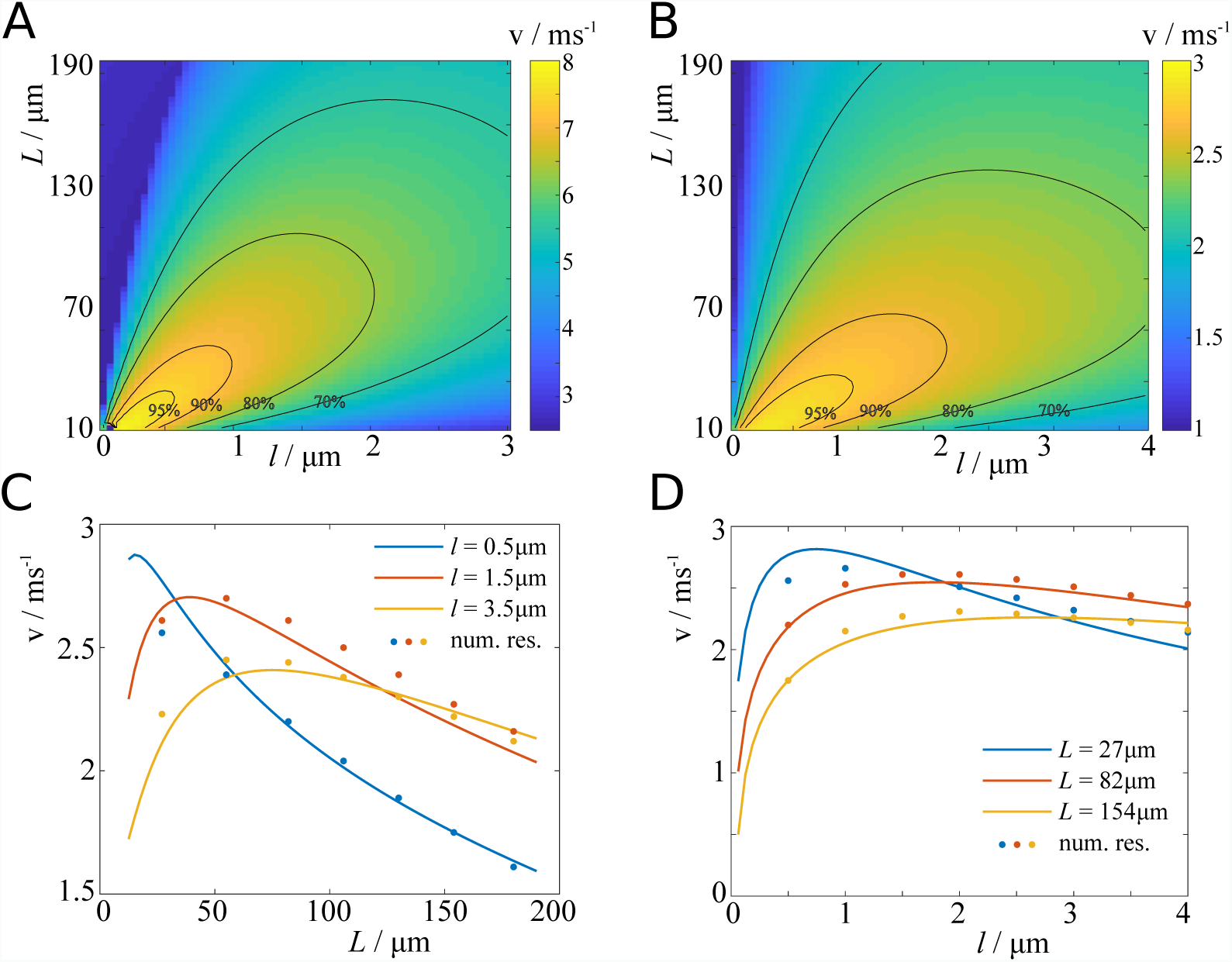
Velocity dependence on node length and internode length. **A**: Propagation velocity plotted against node length and internode length. Contours indicate percentages of maximum velocity. (Scenario D with standard parameters.) **B**: Same as **A**, with fitted parameters. **C**: Propagation velocity as function of internode length (scenario D with fitted parameters), and comparison with numerical results from biophysical model. **D**: Propagation velocity as function of node length, and comparison with the model by Arancibia-Carcamo *et al.* [24].

We find that the propagation velocity varies relatively little with changes in the nodal and internodal length. For scenario D with standard parameters, we find that velocities across the investigated range of parameters are above 70% of the maximum, and for the parameters fitted to the biophysical model the sensitivity is even less. Interestingly, we find that decreasing node length and internode length simultaneously, the velocity increases steadily.

In Fig. 6C and D we show cross-sections of Fig. 6B, and compare these results with the numerical results from the the cortex model used in [24]. There is a good agreement between our model and the biophysical model, with the biggest discrepancies occuring at short node and internode lengths. We assume that these discrepancies arise due to the fact that the biophysical model only uses 50 nodes, whereas we consider *N* = 1000 nodes to determine the velocity. In the Methods section, we show that reducing the number of nodes significantly alters the results at short node and internode lengths (Fig. 13).

#### Myelin thickness

The relative thickness of the myelin layer is given by the g-ratio, which is defined as the ratio of inner to outer radius. Hence, a smaller g-ratio indicates a relatively thicker layer of myelin around the axon. In humans, the g-ratio is typically 0.6−0.7, although it is also known to correlate with the axon diameter [54]. In our mathematical framework, the g-ratio affects the electrotonic length constant *λ* of the internodes, which scales with 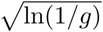. A classical assumption is that the propagation velocity scales in the same manner [1]. Our results suggest (see Fig. 7A) that the velocity depends more strongly on the g-ratio. We therefore generalised this relationship to *v* = *κ*(ln(1/*g*))^*α*^, and find (fitting both *κ* and *α*) our results best match *α* = 0.68 (scenario D with fitted parameters). However, the fitted coefficient *α* also depends on the ratio of internode length and node length, *L/l*. We find that *α* increases monotonically with this ratio (see Fig. 7), and approaches zero when *L/l* approaches zero. The latter represents the case of an unmyelinated axon.

**Fig 7.**
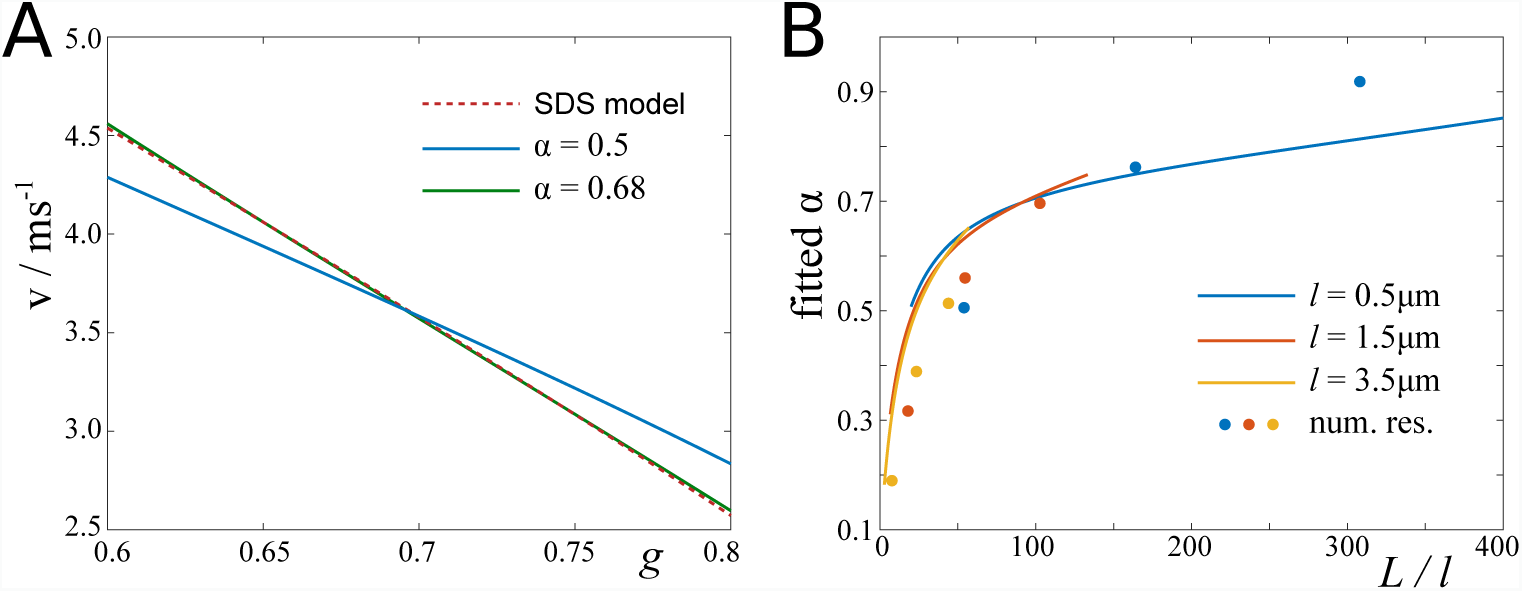
Relative propagation velocity as function of g-ratio. **A**: Result of our spike-diffuse-spike model, and *v* = *κ*(ln(1/*g*))^*α*^ fitted to this result (first with *α* = 0.5 fixed, and then with *κ* and *α* fitted). **B**: Fitted *α* changes with the ratio of internode length to node length in the spike-diffuse spike model (lines), and in the Arancibia-Carcamo model (dots). Parameters: fitted parameters (see Table 1 in Methods section).

In Fig. 8 we present two-parameter plots of the velocity as function of the g-ratio and axon diameter (Fig. 8A), and g-ratio and fibre diameter (Fig. 8B). If the axon diameter is held constant, the velocity increases monotonously with decreasing g-ratio. However, if the fibre diameter is held constant, then the velocity saturates at around *g* = 0.5, because decreasing *g* at constant fibre diameter means decreasing the axon diameter.

**Fig 8.**
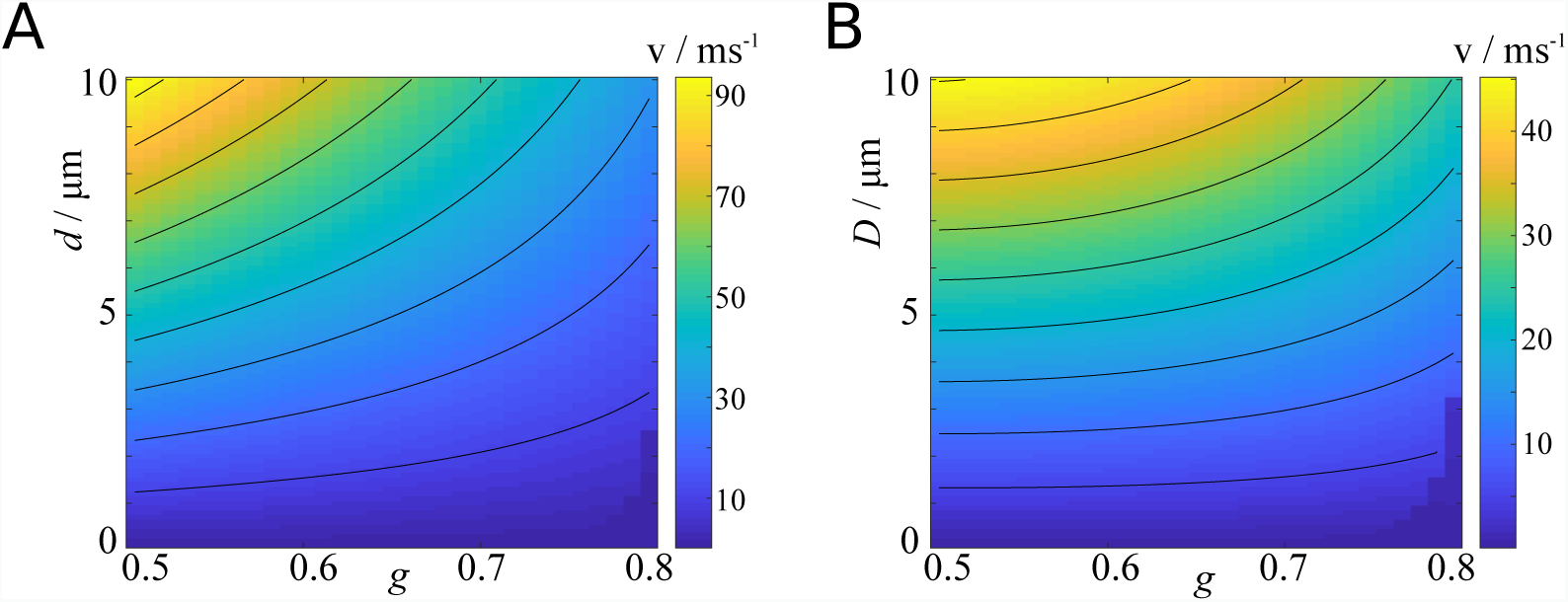
Effect of diameter and g-ratio on propagation velocity. **A**: Velocity plotted against g-ratio and axon diameter. **B**: Velocity plotted against g-ratio and fibre diameter.

#### Ephaptic coupling and entrainment

We demonstrate here that it is possible to study the effects of ephaptic coupling on action potential propagation within our framework. We choose two axonal fibres as a simple test case, but more complicated scenarios could also be considered using our analytical approach. Ephaptic coupling occurs due to the resistance and finite size of the extra-cellular space. We follow Reutskiy *et al.* [31] in considering the axonal fibres being embedded in a finite sized extra-cellular medium (the space between the axons within an axonal fibre bundle). The resulting cable equation for the *n*^*th*^ axon reads

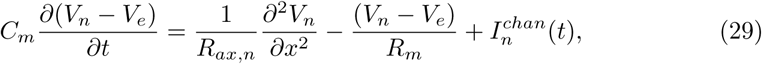

with *V*_*e*_ being the potential of the extra-cellular medium. In the Methods section we describe how to obtain solutions to this set of equations.

We explore solutions to Eq. (29) in a number of ways, which are graphically represented in Fig. 9. We focus on sodium currents as described by scenario D with standard parameters. First, we study how the coupling could lead to entrainment, i.e. synchronisation of action potentials. To this end, we compare the time courses of *V*_1_(*t*) and *V*_2_(*t*) in a pair of axons, where an action potential is emitted in the first axon at *t* = 0, and in the second axon at *t* = ∆*t*. We then compare the *t*_*sp*_ in the neighbouring nodes, and find that for any low threshold values *V*_*thr*_ the difference between the *t*_*sp*_ is less than ∆*t*, meaning the two action potentials are re-synchronising, see Fig 9A. Next, we asked how the coupling affects the speed of two entrained action potentials. Now we set ∆*t* = 0, in which case *V*_1_(*t*) = *V*_2_(*t*). We compare the depolarisation curves of the simultanously active axons with when only one axon is active, and find that the voltages rise more slowly if two action potentials are present, thus increasing *t*_*sp*_ and decreasing the speed of the two action potentials, see Fig 9B. Thirdly, we considered the case when there is an action potential only in one axon, and computed the voltage in the second, passive axon. We find that the neighbouring axon undergoes a brief spell of hyperpolarisation, with a half-width shorter than that of the action potential. This hyperpolarisation explains why synchronous or near-synchronous pairs of action potentials travel at considerably smaller velocities than single action potentials. The hyperpolarisation is followed by weaker depolarisation.

**Fig 9.**
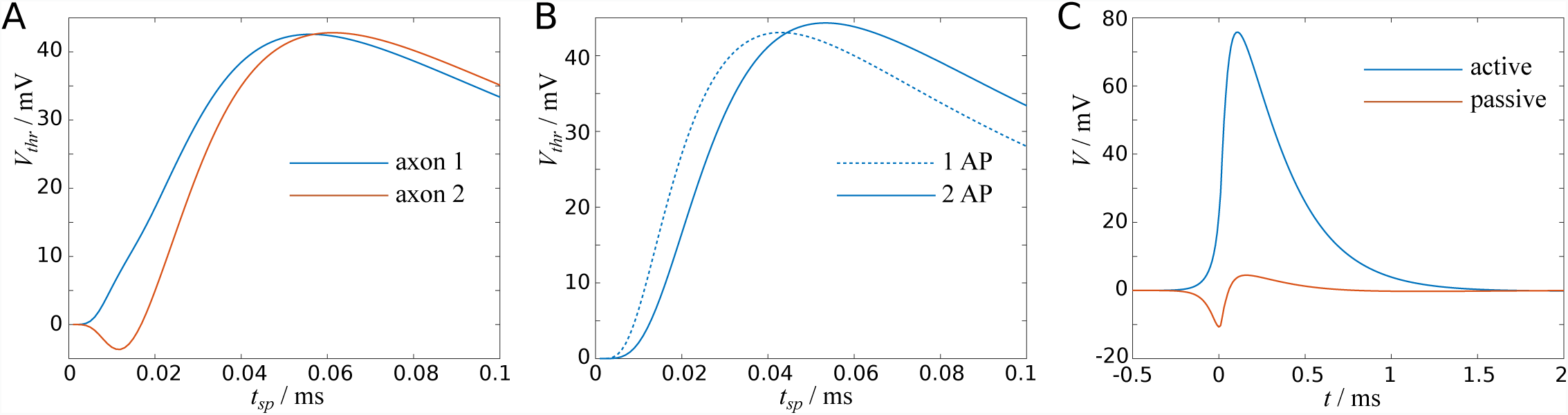
Ephaptic coupling reduces AP speed and leads to AP synchronisation. **A**: Depolarisation curves for a pair of action potentials with initial offset of 0.02ms converge, reducing the time difference between action potentials. **B**: Depolarisation of a synchronous pair of action potentials is slower than for a single action potential. **C**: An action potential induces initial hyperpolarisation and subsequent depolarisation in an inactive neighbouring axon.

## Discussion

We have developed an analytic framework for the investigation of action potential propagation based on simplified ion currents. Instead of modelling the detailed dynamics of the ion channels and its resulting transmembrane currents, we have adopted a simpler notion by which a threshold value defines the critical voltage for the ion current release. Below that threshold value the membrane dynamics is passive, and once the threshold value is reached the ion current is released in a prescribed fashion regardless of the exact time-dependence of the voltage before or after. We studied four different scenarios, of which the simplest was described by a delta-function representing immediate and instantaneous current release. The three other scenario incorporated delays in different ways, from a shift of the delta function to exponential currents and, lastly, combinations thereof. The latter seemed most appropriate considering experimental results.

The simplified description of the ion currents permitted the use of analytical methods to derive an implicit relationship between model parameters and the time the ion current would depolarise a neighbouring node up to threshold value. This involved the solution of the convolution integral of the ion current with the Green’s function of the passive cable equation. From the length of nodes and internodes and the time to threshold value between two consecutive nodes (*t*_*sp*_) resulted the velocity of the action potential.

We only obtained an implicit relationship between the threshold value *V*_*thr*_ and the parameter *t*_*sp*_, which needed to be solved for *t*_*sp*_ using root-finding procedures. However, in comparison to full numerical simulations, our scheme still confers a computational advantage, as the computation time is about three orders of magnitude faster than in the biophysical model by Arancibia-Carcamo *et al.* [24]. In the Methods section we have shown that one can achieve a good approximation by linearising the rising phase of the depolarisation curve. We did not explore this linearisation further, but in future work it might serve as a simple return-map scheme for action potential propagation, in which parameter heterogeneities along the axon could be explored.

We used our scheme to study the shape of action potentials, and we found that the ion currents released at multiple nearby nodes contribute to the shape and amplitude of an action potential. This demonstrates that action potential propagation is a collective process, during which individual nodes replenish the current amplitude without being critical to the success or failure of action potential propagation. Specifically, the rising phase of an action potential is mostly determined by input currents released at backward nodes, whereas the falling phase is determined more prominently by forward nodes (cf. Fig. 4).

Our scheme allowed us to perform a detailed analysis of the parameter dependence of the propagation velocity. We recovered previous results for the velocity dependence on the axon diameter, which were an approximately linear relationship with the diameter in myelinated axons, and a square root relationship in unmyelinated axons. Although the node and internode length are not accessible to non-invasive imaging methods, we found it pertinent since a previous study [24] looked into this using numerical simulations. Our scheme confirms their results qualitatively and quantitatively, and by performing a more detailed screening of the node length and the internode length revealed that for a wide range the propagation velocity is relatively insensitive to parameter variations.

We also studied the effect of the *g*-ratio on the propagation velocity, which was stronger than previously reported, as we find that the velocity is proportional to (ln(*−g*))^*α*^ with *α* ≈ 0.7, whereas the classical assumption was *α* = 0.5 [1]. Furthermore, we found that *α* depends on the ratio between node length and internode length, which to the best of our knowledge has not been reported before. Intuitively, changing the thickness of the myelin sheath of relatively short internodes has a smaller effect than changing the myelin thickness around long internodes (relative to the node length).

The main results of our spike-diffuse-spike model were compared with the biophysically detailed model recently presented by Arancibia-Carcamo *et al.* [24]. The latter uses the Hodgkin-Huxley framework and models the myelin sheath in detail, including periaxonal space and individual myelin layers. To enable the comparison between the two models, we fitted parameters of our spike-diffuse-spike model to output of the the biophysical model. In spite of the differences in the model setup, we find that the results of the two models agree well.

The framework developed here also allowed us to study the effect of ephaptic coupling between axons on action potential propagation. We found that the coupling leads to the convergence between sufficiently close action potentials, also known as entrainment. It has been hypothesised that the functional role of entrainment is to re-synchronise spikes of source neurons. We also found that ephaptic coupling leads to a decrease in the wave speed of two synchronous action potentials. Since the likelihood of two or more action potentials to synchronise in a fibre bundle increases with the firing rate, we hypothesise that a potential effect could be that delays between neuronal populations increase with their firing rate, and thereby enable them to actively modulate delays. In addition, we examined the temporal voltage profile in a passive axon coupled to an axon transmitting an action potential, which led to a brief spell of hyperpolarisation in the passive axon, and subsequent depolarisation. This prompts the question whether this may modulate delays in tightly packed axon bundles without necessarily synchronising action potentials. The three phenomena we report here were all observed by Katz and Schmitt [55] in pairs of unmyelinated axons. Our results predict that the same phenomena occur in pairs (or bundles) of myelinated axons.

There are certain limitations to the framework presented here. First of all, we calibrated the ion currents with data found in the literature. This ignores detailed ion channel dynamics, and it is an open problem how to best match ion currents produced by voltage-gated dynamics with the phenomenological ion currents used in this study. Secondly, we assumed that the axon is periodically myelinated, with constant g-ratio and diameter along the entire axon. The periodicity ensured that the velocity of an action potential can be readily inferred from the time lag between two consecutive nodes. In an aperiodic medium, the threshold times need to be determined for each node separately, resulting in a framework that is computationally more involved. Here it might prove suitable to exploit the linearised expressions for the membrane potential to achieve a good trade-off between accuracy and computational effort. Heterogeneities in the g-ratio or the axon diameter would be harder to resolve, as the corresponding cable equation and its Green’s function would contain space-dependent parameters. If individual internodes are homogeneous, then one could probably resort to methods used in [36] to deal with (partially) demyelinated internodes. Thirdly, we studied ephaptic coupling between two identical fibres as a test case. Our framework is capable of dealing with axons of different size too, as well as large numbers of axons. In larger axon bundles, however, it might be necessary to compute the ephaptic coupling from the local field potentials, as the lateral distance between axons may no longer allow for the distance-independent coupling we used here. Nevertheless, it would be interesting to extent our framework to realistic axon bundle morphologies, and test if the predictions we make here, i.e. synchronisation of action potentials and concurrent increase in axonal delay, still hold. If yes, then there may also be the possibility that delays are modulated by the firing rates of neuronal populations.

## Methods

### The cable equation

To model action potential propagation along myelinated axons, we consider a hybrid system of active elements coupled by an infinitely long passive cable. The latter represents the myelinated axon and is appropriately described by the cable equation, whereas the active elements represent the nodes of Ranvier whose dynamics are governed by parametrically reduced, phenomenological dynamics.

In general, a myelinated axon can be described by the following cable equation:

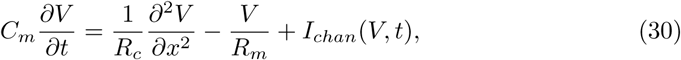

where *V* is the trans-membrane potential, *I*_*chan*_(*V, t*) represents the ionic currents due to the opening of ion channels, and *x* represents the spatial coordinate longitudinal to the cable. *C*_*m*_ and *R*_*m*_ are the capacitance and resistance of myelinated segments of the cable. Multiplying both sides of (30) with *R*_*m*_ yields

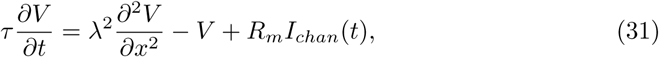

where *τ* = *C*_*m*_*R*_*m*_ and 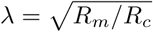 are the time constant and cable constant pertaining to the internodes. All model parameters are listed in Table 1.

### Cable parameters

The capacitance of a cylindrical capacitor (such as a myelin sheath, or the insulating part of a coaxial cable) can be found by considering the following relationship,

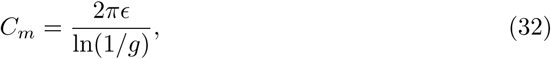

with *g* being the g-ratio, i.e. the ratio between axon diameter and fibre diameter. The parameter ϵ denotes the permittivity of the medium. The radial resistance of the cylinder is given by:

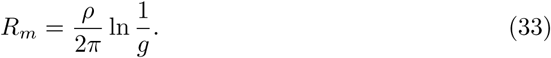

The parameter *ρ* describes the resistivity of the cylindrical medium.

Experimental values for the capacitance and radial resistance of a myelinated axon are reported in Goldman and Albus [20],

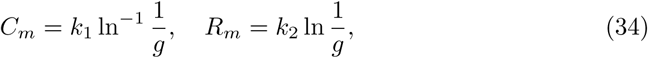

with (taking values from [56] and assuming *g* = 0.8 in the frog)

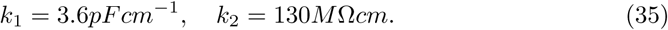

The values for *k*_1_ and *k*_2_ correspond to the following values for permittivity and resistivity:

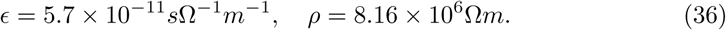

Finally, the axial resistance per unit length along the inner medium of the cylinder is given by

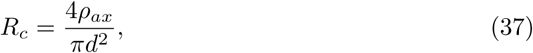

where *ρ*_*ax*_ = 110Ω*cm* [20] is the resistivity of the inner-axonal medium, and *πd*^2^/4 its cross-sectional area.

With these constants at hand, we can now define the parameters of equation (31):

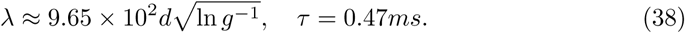

We treat the axonal diameter *d* and the g-ratio *g* as free parameters, and *ρ*_*ax*_, *k*_1_ and *k*_2_ are treated as constants.

### Analytical solution

The inhomogenenous cable equation can be written in compact form:

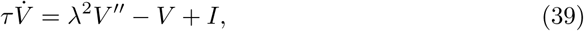

with 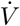 indicating the time derivative of *V*, and *V*″ indicating the second spatial derivative of *V*. Fourier transformation in *x* yields an ordinary differential equation of the form,

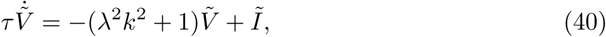

where~indicates the Fourier transformed quantity. The homogeneous part of Eq. (40) has the solution

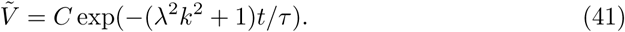

The inhomogeneous solution in *t* can be found by the method of variation of the constant, which yields the following convolution integral in *t*:

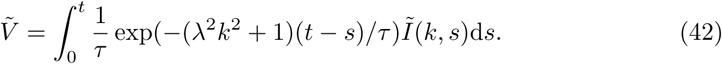

The inverse Fourier transform of Eq. (42) then yields the following double convolution integral in *x* and *t*:

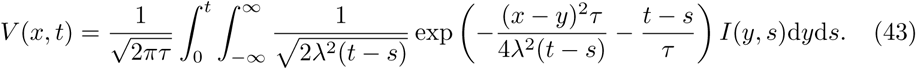

Since we assume the nodes of Ranvier to be discrete sites described by delta functions in *x*, this integral becomes ultimately a convolution integral in time only.

Thus, we can identify the Green’s function of the cable equation (1) as

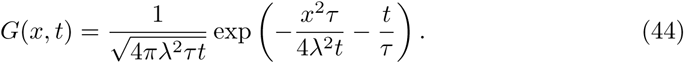

This is Green’s function representing the time evolution of the voltage in a cable due to an instantaneous, normalised input current at distance *x* at time *t* = 0. A graphical representation of *G*(*x, t*) is given in Fig. 10A for various values of *x*.

**Fig 10.**
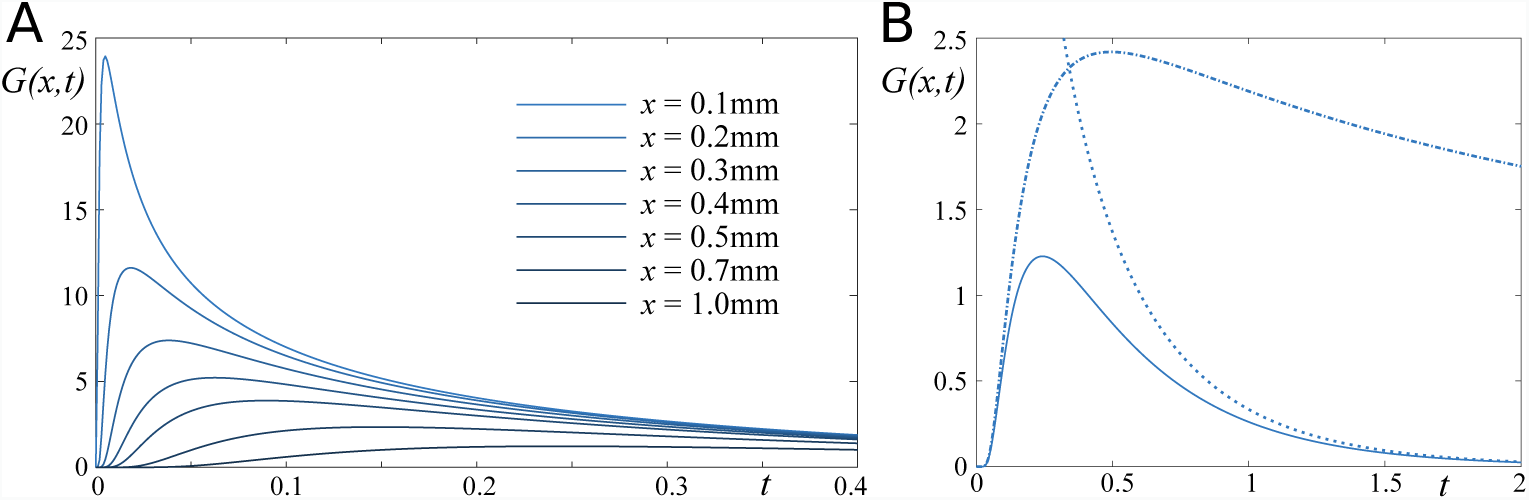
Green’s function of the cable equation. **A**: Green’s function for various distances *x*. **B**: Green’s function for *x* = 1mm, showing the slow (dotted) and fast (dash-dotted) approximation.

We note here that the Green’s function contains *two* time scales. The first is the characteristic time scale of the cable, *τ*, which indicates the voltage decay across the myelin sheeth. The second time constant is *x*^2^*τ*/4*λ*^2^, which is the time it takes exp(−*x*^2^*τ*/4*λ*^2^*t*) to reach 1/e ≈ 0.37. This time depends on all cable parameters, and if *x/λ <* 1 it is significantly faster than *τ*. Hence, if *t ≪ τ*, the cable equation can be approximated by

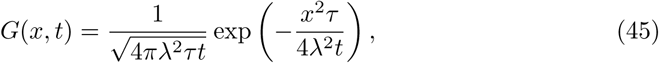

or, conversely, in the limit *t ≫ τ*, it can be approximated by

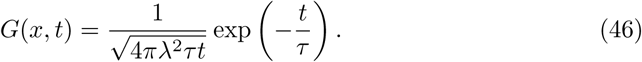

See Fig 10B for a comparison.

### Nodal properties

Like the myelinated parts of the axon, the Ranvier nodes are characterised by their electrophysiological properties through the membrane resistance and membrane capacitance, denoted by *R*_*n*_ and *C*_*n*_, which result in a characteristic length scale *λ*_*n*_ and a characteristic time scale *τ*_*n*_. We use the following values for *R*_*n*_ [20] and *C*_*n*_ [57]:

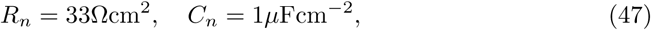

where 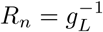, i.e. the inverse leak conductance. With *τ*_*n*_ = *C*_*n*_*R*_*n*_ we obtain a characteristic time of *τ*_*n*_ = 33*µ*s. This value is striking, since typical time constants for neurons at dendrites and the soma range from 10ms to 100ms. This can be explained by the higher density of sodium channels at the nodes of Ranvier than at the soma. As reported in [58], there are approximately 1200 channels per *µ*m^2^ at nodal segments, and only about 2.6 channels per *µ*m^2^ at the soma. Thus, the ratio of ion channel densities between node and soma is nearly 500. We assume here that the conductance scales linearly with the channel density, which is supported by the fact that the membrane resistance is approximately 10kΩcm^2^ at the soma.

### Current influx and separation

The channel current that flows into the axon, *I*_*chan*_(*t*) is counter-balanced by currents flowing axially both ways along the axon, *I*_*cable*_(*t*), and a radial current that flows back out across the membrane of the node, *I*_*node*_:

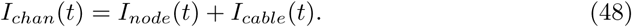

The ratio of currents that pass along the cable and back across the nodal membrane is determined by the respective resistances:

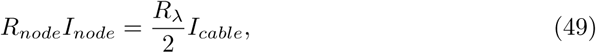

where *R*_*λ*_ is the longitudinal resistance of the axon, defined by *R*_*λ*_ = *R*_*m*_/*λ*. This relationship yields

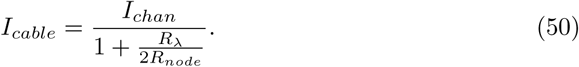

Hence, with the maximum amplitude of the channel current being *I*_0_, the maximum amplitude of current entering the cable is *βI*_0_, where we abbreviate

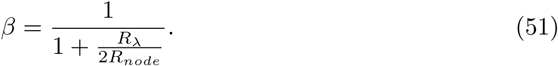

### Approximations and analytical solutions

It is, in general, not possible to find closed-form solutions to the Hodgkin-Huxley model due to the nonlinear dependence of the gating variables on the voltage. We therefore focus here on idealisations of the currents generated by the ion channel dynamics, which is described by a function *I*_*chan*_(*t*).

In mathematical terms, the depolarisation of the neighbouring node is a convolution of the current entering the cable with the solution of the homogeneous cable equation *G*(*x, t*), which describes the propagation of depolarisation along the myelinated segment:

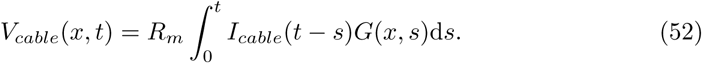

In the following we present the mathematical treatment for the scenarios introduced in the Results section, and we focus here on an input current at a single site.

#### Scenario A - fast current

The (in mathematical terms) simplest scenario is the one in which the ion current is described by the Dirac delta function:

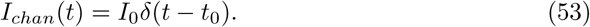

Without loss of generality we set the time of the current, *t*_0_, to zero. The depolarisation along the cable, and specifically at the neighbouring node at distance *x* is then given by the Green’s function of the cable equation itself:

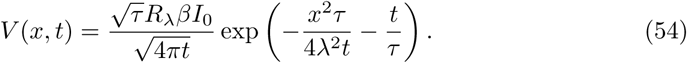

If only one current is injected into the cable, the time *t*_*sp*_ when the threshold value *V*_*thr*_ is reached is given implicitly by

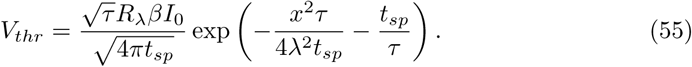

Equation 55 yields an implicit relation for *t*_*sp*_ and the model parameters. There is no obvious way of solving 55 for *t*_*sp*_ explicitly. One can solve it using Newton’s method, and test various parameter dependencies by arc-length continuation. However, we explore the possibility to derive an approximate solution for *t*_*sp*_, and consequently for the axonal propagation speed *v*, by linearisation of (55).

A suitable pivot for the linearisation is the inflection point on the rising branch, i.e. 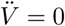 and 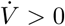. This ensures that the linearisation around this point is accurate up to order 𝒪(*t*^2^), and error terms are of order 𝒪(*t*^3^) and higher. It also provides an unambiguous pivot for the linearisation. Differentiating (54) twice yields

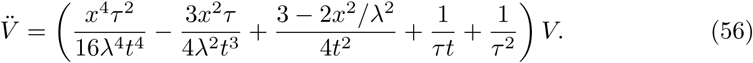

We multiply all terms by *t*^4^ such that the lowest order term in *t* is of order zero. Since *τ* is much larger than the rise time of the depolarisation, we disregard terms of order 𝒪(*t*^3^) and higher. The resulting quadratic equation for the inflection point, *t*_*i*_, yields two positive roots, the smaller of which is

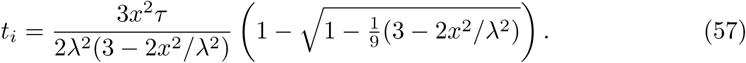

In the limit of *x/λ ≪* 1 we can further simplify this expression to give

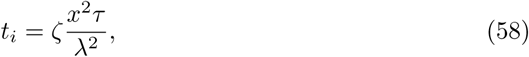

with 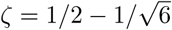. The linear equation for the time-to-spike and the firing threshold is then given by

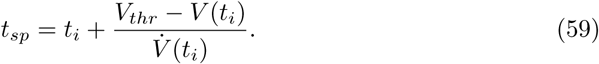

The quantities *V* (*t*_*i*_) and 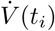 can be approximated to be

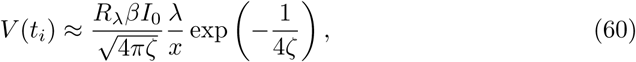

and

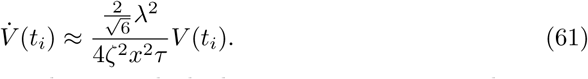

A comparison of the full nonlinear solution with the linear approximation is shown in Fig. 11A.

**Fig 11.**
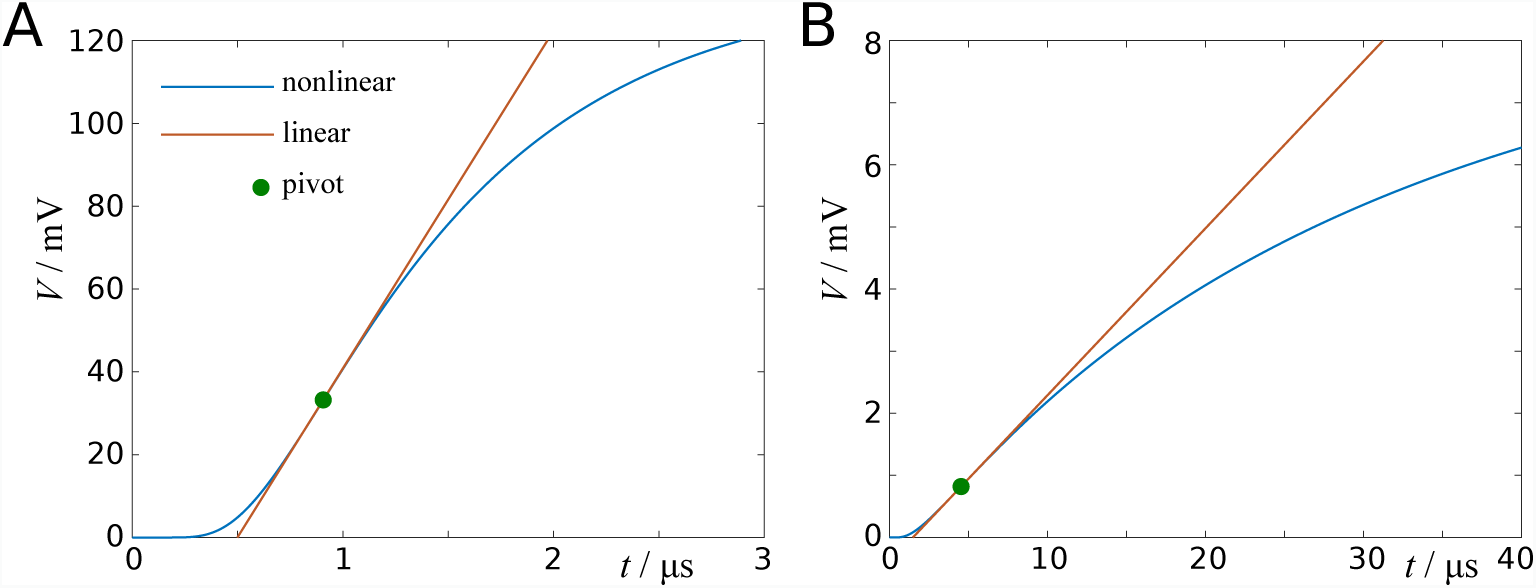
Depolarisation curves and their linear approximation. **A**: Depolarisation curve for instantaneous input current (scenario A). **B**: Depolarisation curve for exponential input current (*τ*_*s*_ = 100*µ*s).

#### Scenario B - delayed fast current

Again we consider a fast current, but one which is emitted with a delay ∆ after the membrane potential has reached the threshold value. If we denote by *t*_0_ the time of the threshold crossing, then the ionic current is given by

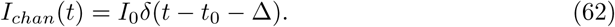

However, by simple linear transformation we may also use *t*_0_ to denote the time of the spike. In this case, a spike will be generated after *t*_*sp*_ + ∆ in the adjacent node, where *t*_*sp*_ is the time to the threshold crossing in the same node, given by equation (55). The speed of a propagating action potential is then given by

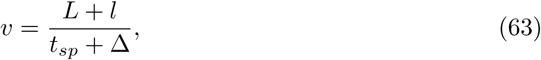

neglecting finite transmission speeds at nodes. In the limit of *t*_*sp*_ → 0 we obtain the result

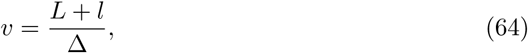

which implies that action potentials can never travel faster than (*L* + *l*)/∆. However, if multiple neighbours are taken into account, the velocity can be faster than this estimate. For example, in Fig. 5A we show results for this scenario with ∆ = 30*µ*s. For an axon diameter of *d* = 1*µ*m (which corresponds to *D* ≈ 1.67*µ*m with *g* = 0.6), we obtain a velocity of about 6m/s, whereas (*L* + *l*)/∆ is approximately 3.3m/s (with *L* = 100*µ*m).

#### Scenario C - exponential current

At this point, we make the assumption that the channel current rises infinitely fast, and drops off exponentially. In mathematical terms, the currents generated by an action potential at a particular node have the following form:

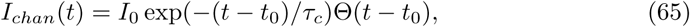

where *I*_0_ denotes the amount of current generated by the channel dynamics, and *t*_0_ denotes the time the spike is generated. The Heaviside step function Θ ensures that *I*_*chan*_(*t*) = 0 for *t < t*_0_. Without loss of generality we set *t*_0_ = 0.

The propagated depolarisation is now given by the convolution of the exponential function with the Green’s function of the cable equation:

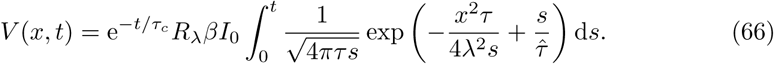

Here we use 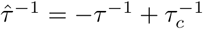. We now briefly sketch how to solve this integral. Disregarding prefactors, the integral to be solved here is of the form

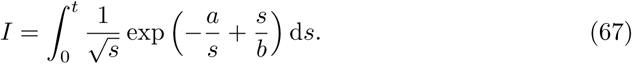

Using the transform 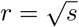 yields

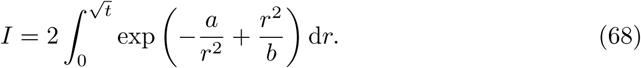

In addition, we define a second integral of the form

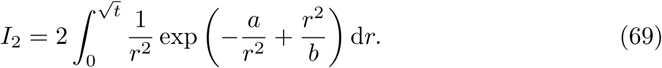

Next, we apply the transform 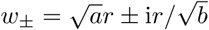 to these two integrals, which yields

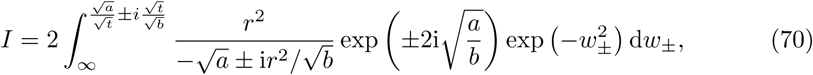

and

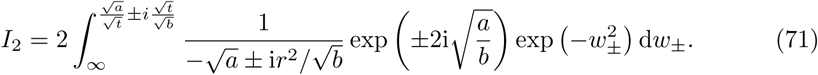

The two integrals can be combined as follows:

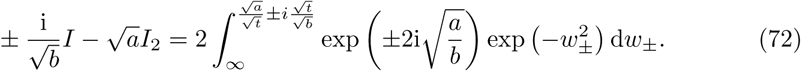

The integral on the right is straightforward to evaluate:

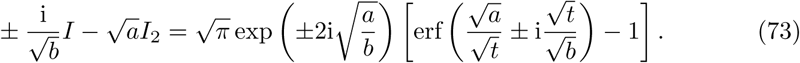

Eliminating *I*_2_ then yields

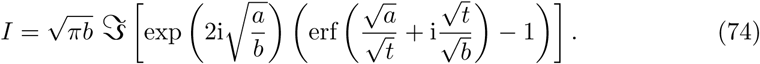

Using the appropriate prefactor and the expressions for *a* and *b*, we finally obtain

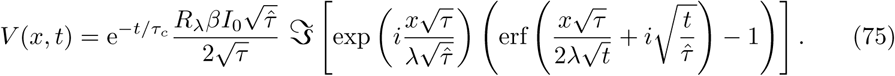

Here, ℑ represents the imaginary part of the argument. The complex argument of the error function arises due to *τ*_*c*_ *< τ*, but this equation also holds if *τ*_*c*_ *> τ* provided that 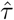 is redefined as 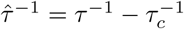.

Once more, we aim to linearise this implicit solution around the inflection point, which in this scenario is identified as 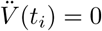. Differentiating *V* (*t*) twice yields

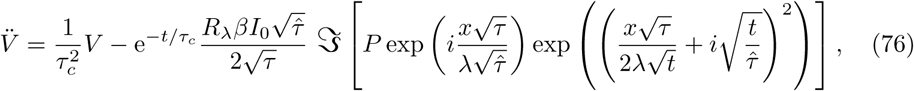

with

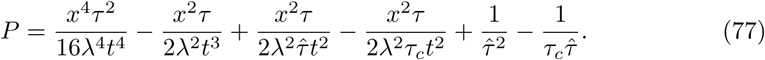

Since the inflection point occurs at small *t*, the terms in *P*(*t*) dominate the curvature of the rising phase of *V*(*t*). Multiplying *P* with *t*^4^ and carrying on terms up to quadratic order then yields the following equation for *t*_*i*_:

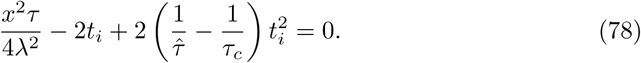

For *τ*_*c*_ *< τ*, this then leads to

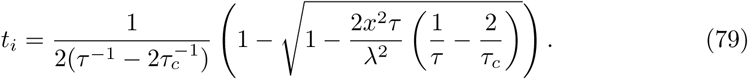

In the limit of *τ*_*c*_ *≪ τ*, this expression reduces to

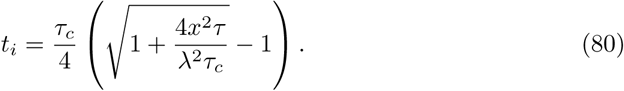

Conversely, if *τ*_*c*_ *> τ*, we find

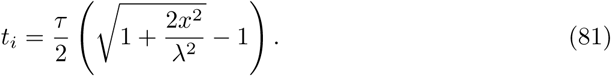

A comparison of the linear approximation with the full nonlinear problem is shown in Fig. 11B.

#### Scenario D - combination of exponentials

Scenario C involved a single exponential function to describe the time course of the channel currents. We now explore more complex time-profiles of channel currents, which can be realised by the sum over *M* exponential time courses with different amplitudes *A*_*s*_ and time constants *τ*_*s*_:

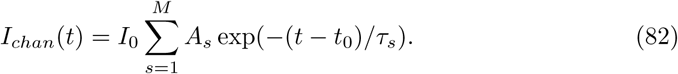

In particular, we consider current profiles of the form

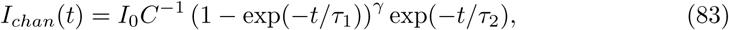

The normalising factor *C* ensures that the maximum value of *I*_*chan*_(*t*) is *I*_0_, which can be determined experimentally. For the sodium current, we use the current density *i*_*Na*_ = 50pA/*µ*m^2^, multiplied by the surface area of the node, throughout the manuscript. This current density yields an amplitude of approximately 100mV for action potentials with standard parameters, although it is twice as high as reported in an experimental study [52]. The reason for the experimental values to be lower might be that for the electrophysiological recordings the axons are severed [59], and ion channels are likely to reorganise and redistribute under such conditions.

Eq. 83 can be recast in the form

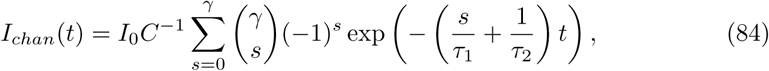

The maximum current is reached at

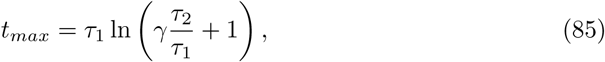

and has the amplitude

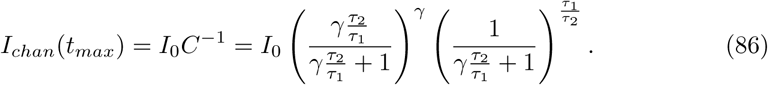

To construct realistic action potentials, we include both sodium and (fast) potassium channels. The sodium gating dynamics of the original Hodgkin Huxley model are governed by a term *m*^3^*h*, where *m* is the activating gating variable, and *h* is the inactivating gating variable. Schwarz *et al.* [60] assume that the dynamics of the resulting ion channel currents can be approximated by

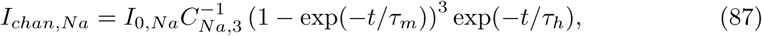

with *C*_*Na*,3_ being the normalisation constant. Baranauskas and Martina [17] presented data that best fit the Hodgkin-Huxley model with *mh*, i.e. a linear relationship with the activating gating variable *m*. In this case, the activation current in our framework reads

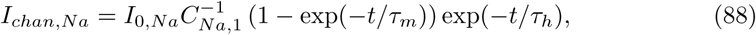

with *C*_*Na*,1_ being the normalisation constant for *γ* = 1. The parameters *τ*_*m*_ and *τ*_*h*_ represent the time scales of the activation and inactivation of the sodium ion channels. Throughout this article we use Eq. (88) to describe the sodium channel dynamics. The time constants are chosen such that the resulting action potential fits best the numerical results for the cortex model in [24], see Fig. 12 for a graphical comparison.

Likewise, we can define the potassium current as follows:

**Fig 12.**
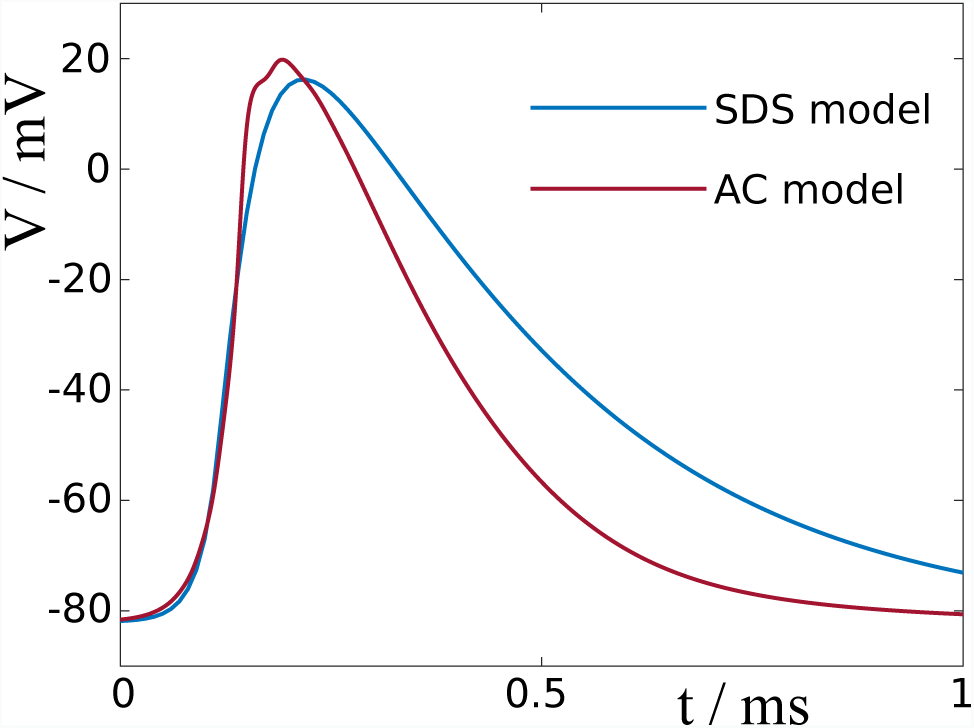
Comparison of action potentials in spike-diffuse-spike model and biophysical model. We chose the time scales *τ*_*m*_ = 20*µ*s and *τ*_*h*_ = 40*µ*s such that the profile, and in particular the rising phase of the action potential in the spike-diffuse-spike model matches well the action potential of the cortical axon model by Arancibia-C`arcamo *et al.* [24].

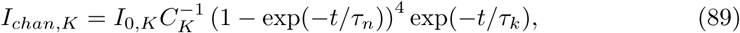

with

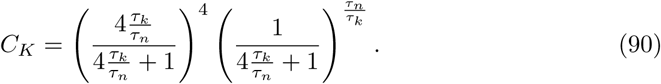

Here, *τ*_*n*_ represents the time scale of the activation of the potassium ion channels. Although there is no inactivating current for potassium in the Hodgkin-Huxley model, we define *τ*_*k*_ as characteristic time with which the potassium current decays. The time constants are voltage-dependent [60], but for simplicity we assume here that they remain constant throughout the formation of the action potential. The peak current density *i*_*K*_ = 3.75pA/*µ*m^2^ is 7.5% of *i*_*Na*_, a ratio we derive from the sodium and potassium conductances used for myelinated axons in [61] (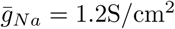 and 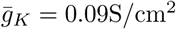).

Finally, denoting the solution to an exponential input current with time constant *τ*_*s*_ by

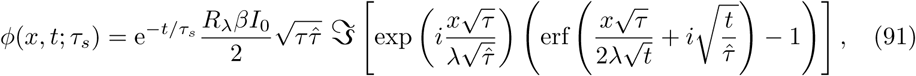

allows us to express the solution as combinations of exponential currents by

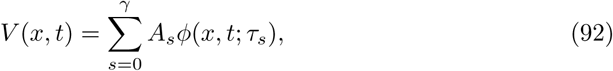

with

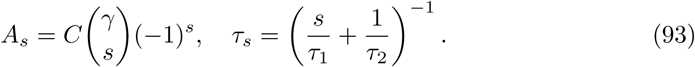

Once more we seek to identify the inflection point, i.e. where 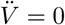. The different time scales *τ*_*s*_ make it difficult to find a closed-form solution, as the ones we found for the previous scenarios. However, we find that a suitable approximation for the inflection point is

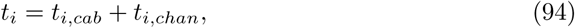

where *t*_*i,cab*_ is the inflection point of the Green’s function of the cable equation in the limit of *x/λ ≫* 1, and *t*_*i,chan*_ is the inflection point of the rising phase of the ion current. *t*_*i,cab*_ can be derived from Eq. 57,

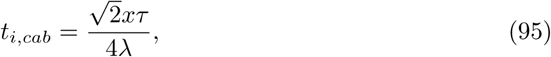

and *t*_*i,chan*_ is found to be

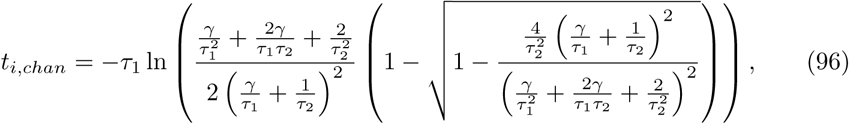

with *γ*, *τ*_1_, and *τ*_2_ as in Eq. 83.

### Influence of distant nodes

Action potentials are driven by the ionic currents generated at multiple nodes along the axon. Due to the linear nature of the cable equation, the effect of multiple input currents can be described by linear superposition:

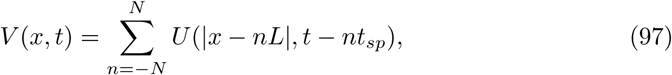

where *U* is the r.h.s. of the respective scenario considered, i.e. *U*(*x, t*) describes the depolarisation due to the current at a nearby node. To keep with our previous definition, time is defined by setting *t* = 0 when the neighbouring node depolarises. The relationship between the firing threshold *V*_*thr*_ and the time-to-spike *t*_*sp*_ is therefore given by

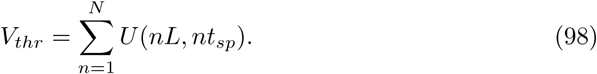

The effect of distant nodes is dampened by the fact that in addition to passing along myelinated segments, currents from distant sources also pass by unmyelinated nodes, and thereby further lose amplitude. Because the distance between two points on the cable is given by *L/λ* in the cable equation, the added distance due to a node with finite length is *l/λ*_*n*_. Therefore, the physical distance between two consecutive nodes is *L* + *l*, and their electrotonic distance is *L* + (*λ/λ*_*n*_)*l* in units of *λ*. This leads to the updated equation for the membrane potential, Eq (9) in the Results section.

As we have shown in Fig. 4, the formation of an action potential is a collective process that incorporates ion channel currents from multiple nearby nodes. Throughout the manuscript we set *N* = 10^3^ to ensure all currents are incorporated, although for the standard parameters *N* = 20 would produce very similar results. However, as we show in Fig. 13, reducing *N* can lead to a considerable reduction of the propagation velocity at short internode lengths.

**Fig 13.**
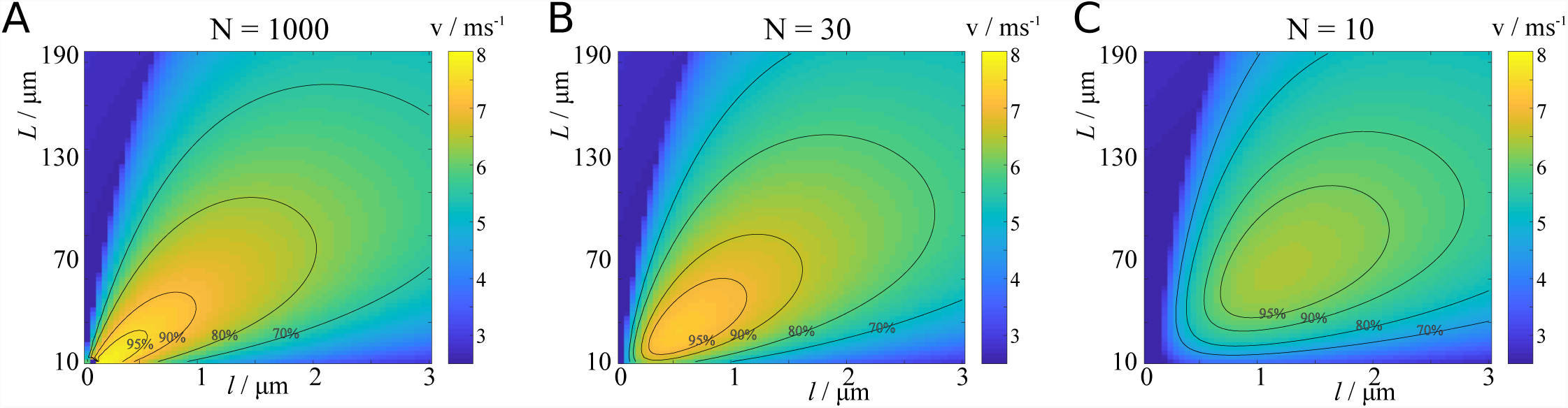
Effect of number of nearest nodes on velocity. We demonstrate here that considering only a small number of nodes can lead to considerable discrepancies in the computed velocity at small node and internode lengths.

This framework allows us to describe unmyelinated axons as well. Since the internode length is zero in this case, the node length *l* is now an arbitrary discretisation of the axon. The membrane potential is now described by

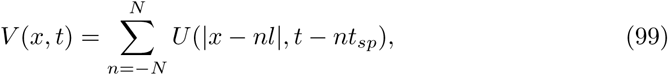

where the length constant *λ* in *U* needs to be replaced by a length constant 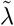 that characterises the electrotonic length of the unmyelinated axon. We introduce a parameter *ρ* that describes the channel density of the unmyelinated axon relative to the channel density of a node of Ranvier. We assume that the conductivity of the axonal membrane scales linearly with the channel density, which implies that the electrotonic length constant of an unmyelinated axon is 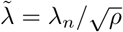, and its time constant is 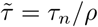. The velocity of an action potential is now defined as *v* = *l/t*_*sp*_.

In addition to the correction terms introduced in Eq (9), we also investigate delays that occur at the nodes due to finite transmission speeds. We assume that action potentials travel with velocities *v* determined by Eq (9) along myelinated segments, and with velocities *v*_*n*_ inferred from Eq (99) at nodes. The corrected velocity is then given by Eq (12) in the Results section.

### Ephaptic coupling and entrainment

Here we explain how to solve Eq (29) with non-zero extra-cellular potential. The potential between intra-cellular medium and extra-cellular medium is *P*_*n*_ = *V*_*n*_ − *V*_*e*_, which determines the channel dynamics. It follows from the electric decoupling of the fibre bundle from the external medium that the sum of longitudinal currents within the fibre bundle is zero [31]:

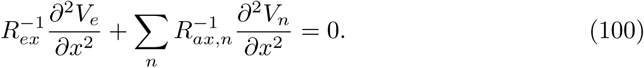

*R*_*ex*_ denotes the axial resistance of the extra-cellular medium, which depends inversely on its cross-sectional area. As a result, we obtain the cable equation in terms of *P*_*n*_:

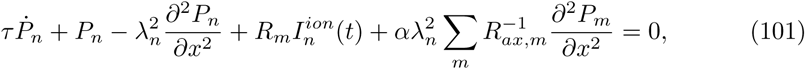

where *α* is the coupling parameter:

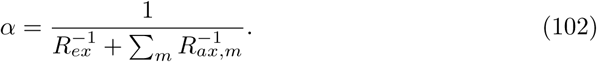

This is a general result, but in the following we focus on two fibres.

Since these equations are linear, they can be decoupled (using orthogonalisation) into

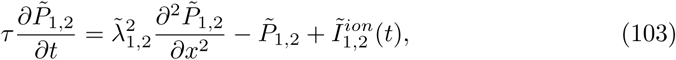

with 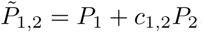, 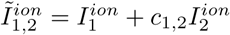, and 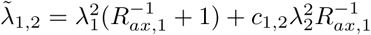, where

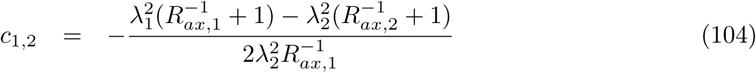

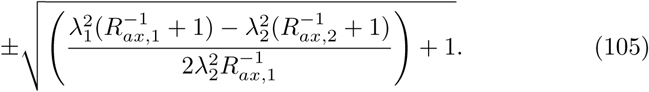

In the case of identical axons, this expression simplifies to *c*_1,2_ = ± 1. These equations can be solved as above, and the solutions of the coupled equations can be recovered using 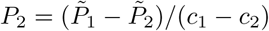 and 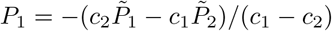.

### Fitting parameters to biophysical model

In order to compare the spike-diffuse-spike model with the biophysical model presented in [24], we generate data points using the biophysical model for the parameters reported therein for the cortex model, and fit our model parameters to these data points. We define a grid of 3 × 3 data points in *L − l*-space for *L* = 27*µ*m, *L* = 82*µ*m and *L* = 152*µ*m, and *l* = 0.5*µ*m, *l* = 1.5*µ*m and *l* = 3.5*µ*m. On this grid we determine the action potential velocity of the biophysical model, which is treated as data for the fitting procedure. Next, we use the least squares curve fit as implemented in MATLAB to fit the following eight parameters of the spike-diffuse-spike model to the data: *λ*, *τ*, *λ*_*n*_, *τ*_*n*_, *τ*_*m*_, *τ*_*h*_, *I*_0_, and *V*_*thr*_. The reason why we use this fitting procedure is that there is no direct correspondence between our model and the biophysical model. The latter implements a Hodgkin-Huxley formalism, as well as a detailed model of the myelin sheath that models each membrane individually and includes periaxonal space. We used the code made available on github by the authors of [24].

## Acknowledgments

HS was supported by the German Research Foundation (DFG Priority Program 2041 ‘Computational Connectomics’, awarded to TRK)

